# Denaturation-sensitive BRET probes enable intracellular thermal shift measurements across diverse protein targets and biomolecular complexes

**DOI:** 10.64898/2026.11.10.765695

**Authors:** James D. Vasta, Ani Michaud, Michael T. Beck, Inhong Hwang, Ashley Schwarzenstein, Cesear R. Corona, Paula Sequeira, Cara Happe, Martin P. Schwalm, Bryan Gehrke, Kendra Nealey, Steven Russell, Bryn Mikulsky, Kaitlin K. Dunn-Hoffman, Cameron Damgaard, Min Zhou, Sophia Berry, Laura Usselmann, Calum Forbes, Mercedes Vazquez-Chantada, Thomas B. Lentz, Rebecca L. Johnson, Michael R. Slater, Cassidy Ames, Magdalena M. Szewczyk, Dalia Barsyte-Lovejoy, Wenhui Zhou, Mei Cong, Poncho L. Meisenheimer, Joseph Shaw, Matthew B. Robers

## Abstract

Thermal shift assays have revolutionized drug discovery, but can be limited in sensitivity and scalability. Combining the unique attributes of differential scanning fluorimetry (DSF) and cellular thermal shift (CETSA), we report the methodological framework for the bioluminescence resonance energy transfer shift assay (BRETSA). BRETSA utilizes denaturation sensitive BRET probes, in combination with luciferase-tagged target proteins, to track and longitudinally record thermal shift in living cells. Denaturation profiles were successfully measured for over 500 globular proteins, spanning 24 target families in diverse cellular compartments. Among these, target engagement was confirmed for over 200 proteins; including validated drug targets as well as understudied members of the kinase subfamily. Leveraging orthosteric, allosteric, and covalent ligands in potency studies, this method offers improved sensitivity and scalability compared to other thermal shift methods. We extend the utility of BRETSA to interrogate target engagement at biomolecular complexes and query cooperative binding for molecular glues and emerging tricomplex drug modalities.

## Introduction

Measuring drug–target interactions in the native cellular environment remains a central challenge in early drug discovery. Biochemical binding assays using purified proteins, though quantitative and scalable, cannot capture the influence of membrane permeability, intracellular concentration gradients, endogenous protein complexes, and post-translational modifications on compound pharmacology. Moreover, the reliance on purified proteins as the substrate for both binding assays and biophysical screening methods restricts chemical biology efforts to the fraction of the proteome amenable to recombinant expression and purification. This represents a significant barrier, as fewer than 15% of human proteins yield soluble, full-length protein in standard expression systems^1^. It is therefore not surprising that the majority of the proteome remains understudied, with a minor fraction successfully targeted for drug discovery^2, 3^. The cellular thermal shift assay (CETSA) was developed to address this gap by exploiting an elegant biophysical principle: ligand binding thermodynamically stabilizes a target protein against thermal denaturation, and this stabilization can be detected in intact cells by measuring the fraction of target that remains soluble after heat challenge^4^. CETSA and its proteome-wide extension, thermal proteome profiling (TPP), have largely revolutionized target validation and off-target identification^5^. High-throughput adaptations using split NanoLuc reporters (SplitLuc CETSA/BiTSA) further extended the approach to 384- and 1536-well microplate formats, enabling screening-compatible throughput^6-10^.

Despite these advances, all CETSA-based methods share a fundamental mechanistic dependence on phase separation as the detection event. Following thermal denaturation, the unfolded target physically aggregates and partitions into the insoluble fraction to produce a measurable change in signal, generally detected by immunoblot, dual-antibodies, or reporter complementation. This phase-separation requirement introduces several technical considerations. Aggregation kinetics are protein-specific and do not always mirror denaturation thermodynamics^11, 12^, which may distort apparent thermal transitions. The lytic endpoint detection also precludes longitudinal measurements of thermal shift in intact cells, requiring large numbers of samples to establish target melt profiles. Finally, physical separation between soluble and insoluble analytes can be laborious and incompatible with scalable assay design.

Differential scanning fluorimetry (DSF) offers a conceptually distinct approach. Rather than relying on phase separation, DSF detects the denatured state directly using environmentally sensitive fluorescent dyes that entropically decorate exposed hydrophobic surfaces of unfolded proteins^13, 14^. This homogeneous, separation-independent readout provides robust and reproducible thermal shift measurements that have become a standard tool in chemical biology and biochemical drug screening^15^. New adaptations of DSF have vastly extended the range of target compatibility^16^. However, DSF requires micromolar concentrations of purified protein, which excludes the cellular context and also creates an assay wall that limits sensitivity.

If the denaturation-sensitive detection principle of DSF could be rendered target-specific, it would in principle combine the best attributes of both approaches: the direct, separation-independent detection of DSF with the physiological relevance of a cellular assay. Bioluminescence resonance energy transfer (BRET) provides a means to achieve this specificity. In BRET, energy transfer from a bioluminescent donor to a proximal fluorescent acceptor occurs only within a Förster radius of approximately 2 nm, imposing stringent distance and geometry constraints that suppress background signal from non-proximal fluorophores^17^.

The tight distance dependence of BRET supports that even a promiscuous, non-selective acceptor dye, such as an environmentally sensitive probe that decorates thousands of denatured proteins, would produce target-specific signal only from the tagged protein of interest. A BRET-based thermal shift assay, combining a thermostable bioluminescent reporter fused to a target protein with a cell-permeable, denaturation-sensitive acceptor probe, could enable intracellular thermal shift analysis. Cell-based analysis would support use of full-length proteins, often incompatible with DSF analysis. A key advantage would be afforded by the homogeneity and robustness of DSF but with target-specific cellular pharmacology as the output.

Here we describe the development and validation of this approach, termed the BRET Shift Assay (BRETSA), including the first complete description of the assay methodology and analytical framework for engagement scoring. We demonstrate broad proteome compatibility across more than 500 target proteins and multiple subcellular compartments, rank-order agreement with orthogonal target engagement methods, and sensitivity advantages over both DSF and aggregation-based cellular assays. We extend the applicability of BRETSA to broad-spectrum interrogation of the human kinome and the ability to measure apparent thermodynamic cooperativity at drug-induced ternary complexes in living cells.

## Results and Discussion

### A BRET-based thermal shift assay for measuring target engagement in living cells

A BRET-based thermal shift assay, leveraging a denaturation-sensitive acceptor probe, could enable intracellular thermal shift analysis that is independent of phase separation **[Fig. 1a].** Furthermore, because BRET can operate with sub-nM concentrations of analyte^17^, BRETSA analysis could offer sensitivity far beyond that of biochemical DSF. To extend the DSF principle into intact living cells, the denaturation-sensitive acceptor probe must permeate cell membranes with minimal toxicity, while preserving membrane integrity. The probe must also report ligand binding with a signal window sufficient to resolve compound effects across diverse targets. A series of denaturation-sensitive acceptor probes were evaluated toward this goal. BRETSA Probe 01 (Promega) is offered specifically for live-cell BRETSA studies and was the focus of this investigation. In addition, SYPRO dyes were included, as these reagents have been applied widely to cell-free DSF studies^14^, and in a limited manner to BRET-based thermal shift in cell extracts^18^. Over its recommended concentration range (up to a 2X dose), BRETSA Probe 01 had negligible cytotoxicity or impacts on membrane integrity **[Fig. 1b, Extended Data Fig. 1a].** In contrast, titration of SYPRO dyes over a dose range commonly used in DSF experiments^14^ **[Fig. 1b, Extended Data Fig. 1a]** resulted in dose- and time-dependent cytotoxicity, with a temporal pattern similar to that of a lytic dose of digitonin. Accordingly, sub-toxic concentrations of the SYPRO dyes were identified and evaluated in live-cell BRET experiments using a series of sentinel targets as proof-of-concept. As sentinel targets, we selected 10 globular, full-length, proteins spanning 3 intracellular contexts and 8 protein families that would stringently test the assay’s capabilities **[Table 1]**. The sentinel set included prototypical thermal shift targets such as MAPK14 (p38α, 41.3 kDa), and also large, membrane-anchored receptor tyrosine kinases MET (156 kDa) and DDR1 (101 kDa), which are often considered more challenging for thermal shift in their full-length form, owing to integral membrane partitioning and large multidomain characteristics.

**Figure 1.**
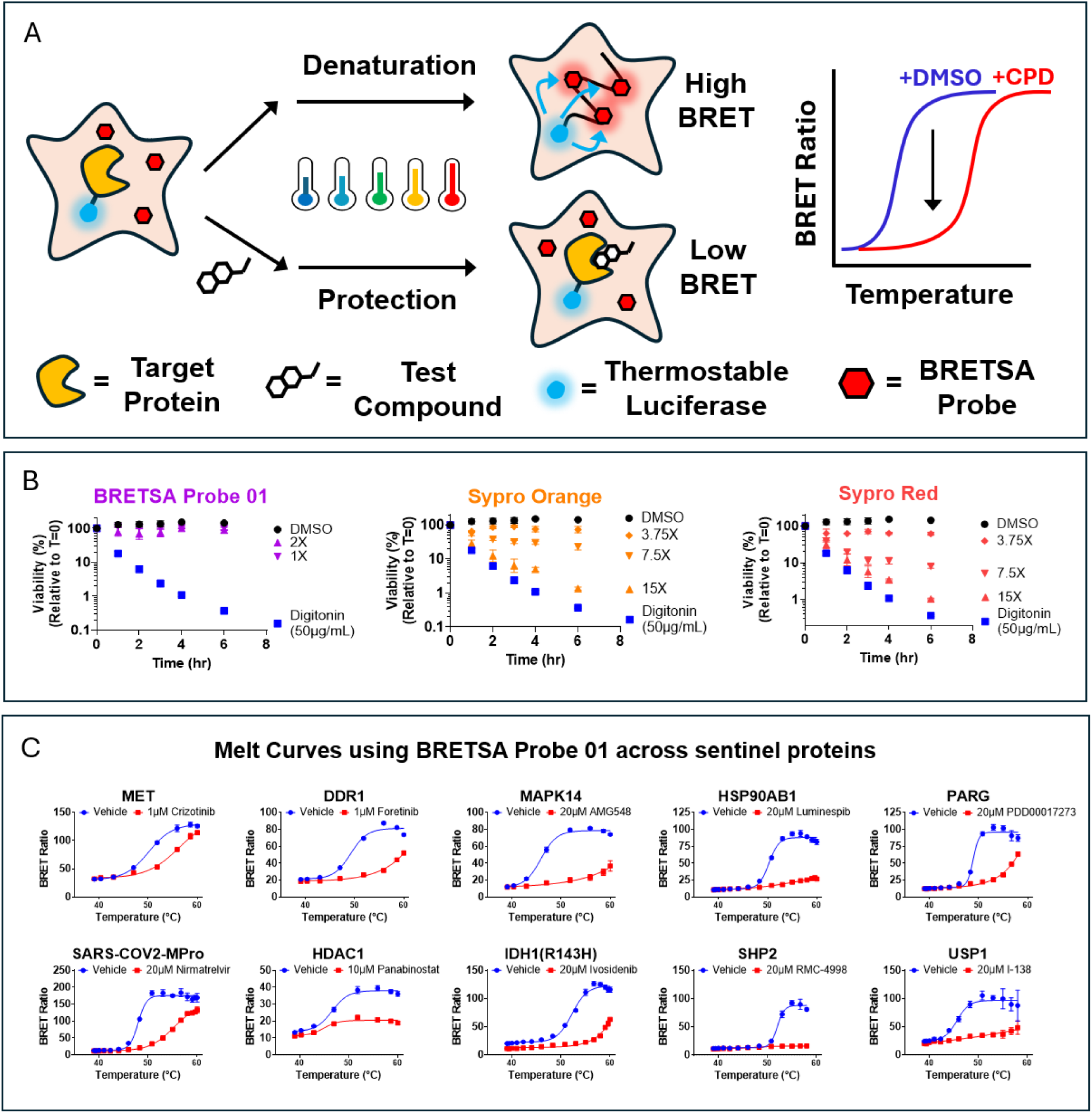
Identifying an ideal probe for the BRET Shift Assay (BRETSA). (A) Schematic overview of the BRETSA approach. A protein of interest fused to a thermostable luciferase is expressed inside living cells. A cell permeable denaturation sensitive acceptor probe (BRETSA probe) is added to the cells. Cells are exposed to a thermal challenge such as a temperature gradient in the presence and absence of a test compound. BRETSA probes that bind to denatured target protein become excited via bioluminescence resonance energy transfer from the pendant thermostable luciferase, resulting in a high BRET signal. Test compounds that bind to and modulate the stability of the target protein will result in a change in the BRET signal, which can be visualized as a shift in a BRET-based thermal stability curve. (B) HEK-293 cytotoxicity curves for BRETSA Probe 01, Sypro Orange, or Sypro Red. Data are representative of experiments performed twice, where individual data points are the mean (± SD) of 4–8 technical replicates. (C) Example BRETSA melt curves for 10 target protein-luciferase fusions expressed inside live HEK-293 cells and treated with a saturating dose of a known cell permeable ligand. Data are representative of experiments performed twice, where individual data points are the mean (± SD) of 3–4 technical replicates. All targets used a dose of 1X BRETSA Probe 01, except for MET kinase, which used a 2X dose.

**Table 1:**
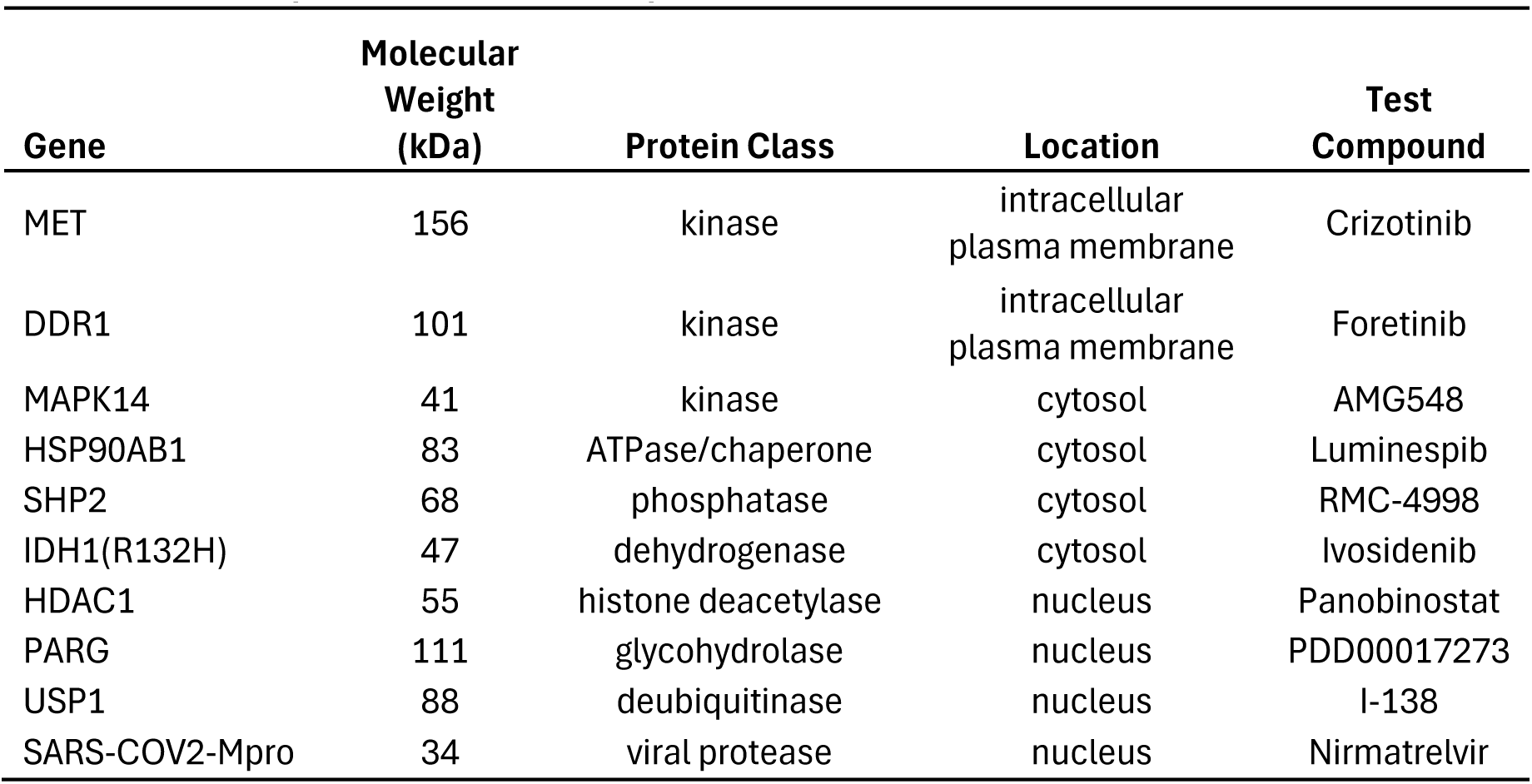
Sentinel protein set for BRETSA probe selection.

To evaluate BRET and compound effects in cells, luciferase fusions were introduced on each gene and ectopically expressed in HEK-293 cells. The reporter was fused to the N- or C-terminus on each protein to ensure proximity to the ligand binding domain, thus maximizing the coupling between ligand-induced stabilization of the binding domain and suppression of the BRET signal. Cells were then subjected to the assay workflow described in **Fig. 1a**. In this workflow, an extracellular NanoLuc inhibitor was included to conditionally measure a luminescent signal in intact cells, while gating luminescence signal from compromised cells^19^. In intact cells, BRETSA Probe 01 detected thermal protection (i.e., stabilization) with a representative test compound for each sentinel target **[Fig. 1c, Extended Data Fig. 1b]**. At the highest sub-toxic concentration within the conventional dose range, SYPRO dyes showed modest BRET signals for the prototypical thermal shift target MAPK14, but generated relatively low signal-to-background for many other sentinel targets including the membrane-associated targets MET and DDR1 **[Extended Data Fig. 1c]**. To facilitate comparison of the performance between the various thermal shift probes, we calculated the amplitude of BRET inhibition for the compound effect (isothermal engagement window, see methods) as a function of temperature for each target/compound pair **[Extended Data Fig.1c]**. Categorically, BRETSA Probe 01 provided the largest amplitude of compound effect (mean 4.1 ± 1.6-fold). Moreover, the SYPRO dyes caused spurious dose-dependent inhibition of the luminescent donor signal in live cells **[Extended data Fig. 1d].** Based upon the superior performance of BRETSA Probe 01, reflected by the observed cytotoxicity, weak signal-to-background, and compromised luminescence observed with the SYPRO dyes, BRETSA Probe 01 was selected for an expanded study across intracellular targets and modalities.

### Broad proteome compatibility across target classes and cellular compartments

For cellular denaturation analysis over a range of diverse intracellular proteins, the luminescent reporter requires two critical features. First, the reporter must be sufficiently thermostable and remain folded while the target protein denatures. NanoLuc, for example, is thermolabile in the 60-70°C range, representing a potential limitation for highly thermostable proteins^7^. Furthermore, luciferases such as NanoLuc are often highly active in cellular debris, an issue likely exacerbated by high temperatures^7^. For conditional live-cell analyses, the ideal reporter should not only offer high thermostability, but it must also be sensitive to the extracellular inhibitor that suppresses luciferase signal from compromised cells^19^. StabiLuc was engineered as a distinct thermostable NanoLuc variant (stability > 100°C) that also retains full sensitivity to the extracellular inhibitor **[Fig. 2a and 2b; Extended data Fig. 2a and 2b]**. As a proof of concept, BRETSA was demonstrated using StabiLuc with KRAS(G12C), a protein that is both highly stable in its nucleotide-bound state and known to be further stabilized by covalent switch-II pocket ligands such as sotorasib^20^. Using BRETSA, StabiLuc-KRAS(G12C) demonstrated a T_M,app_ of 63°C and a clear stabilizing effect from sotorasib treatment **[Fig. 2c; Supplementary Data Set 1],** both in excellent agreement with that reported previously^20^.

**Figure 2.**
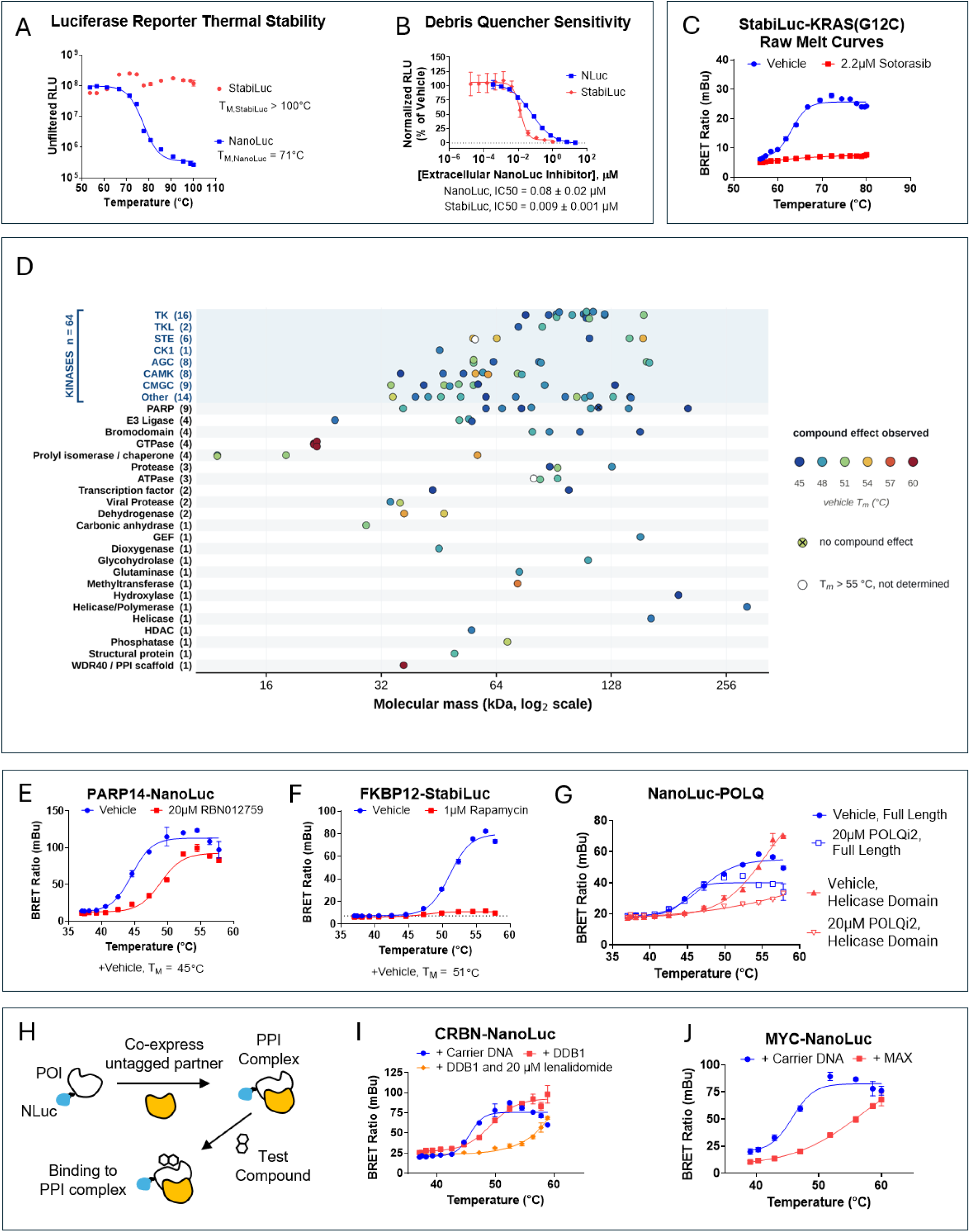
Broad target survey using StabiLuc and NanoLuc reporters. (A) Thermal stability of the NanoLuc and StabiLuc luciferases based upon light output as a function of thermal challenge. Data are representative of two independent experiments, where each data point represents the mean (± SD) of 5 technical replicates. (B) NanoLuc and StabiLuc sensitivity to the extracellular NanoLuc inhibitor (Debris Quencher) as measured by titrating the inhibitor into cells expressing either luciferase after digitonin permeabilization. Data are representative of two independent experiments, where each data point represents the mean (± SD) of 3–4 technical replicates. (C) Representative BRETSA melt curve of StabiLuc-KRAS(G12C) in the presence and absence of sotorasib treatment. Data are representative of two independent experiments where individual data points are the mean (± range) of 2 technical replicates. (D) Graphical summary of the broad target survey representing 114 unique target/ligand interactions across 24 protein families. Coloration of each data point indicates the apparent melting temperature of the protein. Data are summarized from Supplementary Data Set 1 and Extended Data Table 1. (E–G) Example BRETSA melt curves in the presence or absence of test compound (as indicated) for PARP14 (E), FKBP12 (F), and both the full length and helicase domain of POLQ (G). Data are representative of two independent experiments where individual data points are the mean (± range) of 2 technical replicates. Biological replicates are provided in Supplementary Data Set 1. (H) Schematic overview of protein-protein interaction (PPI) assessment and target engagement at a PPI complex using partner protein co-expression studies. (I) Influence of DDB1 co-expression and/or lenalidomide treatment on thermal stability of CRBN-NanoLuc using BRETSA. Data are representative of two independent experiments where individual data points are the mean (± range) of 2 technical replicates. (J) Assessment of the PPI between MAX and MYC-NanoLuc using BRETSA. Data are representative of two independent experiments where individual data points are the mean (± SD) of 3–6 technical replicates.

To evaluate the generality of BRETSA, we assembled a panel of targets (encoding NanoLuc or StabiLuc fusions) spanning diverse protein classes, molecular weights, and subcellular localizations. For each target, the reporter fusion orientation (N- or C-terminal) was selected based on the predicted proximity between the protein terminus and the ligand binding domain, using available structural information and UniProt-annotated domain boundaries^21^. However, for a number of relatively small protein targets (e.g. CypA, CRBN, FKBP12, GRK7, MDM2, VHL, VRK1, and p53(Y220C), both N- and C-terminal tagging was evaluated. Using a standardized 384-well thermal matrix protocol (temperature gradient × compound dose), we generated melt profiles for 114 known globular, full-length target–ligand pairs across 24 distinct families, encompassing kinases, E3 ligases, PARPs, bromodomains, HDACs, deubiquitinases, chaperones, ATPases, GTPases, phosphatases, transcription factors, human and viral proteases, helicases, structural proteins, and various metabolic enzymes **[Fig. 2d; Supplementary Data Set 1].** Validated targets demonstrating a compound effect span molecular weights from ∼12 kDa (FKBP12) to > 200 kDa (PARP14 and POLQ) **[Fig. 2e–g and Supplementary Data Set 1]** and reside in the cytoplasm, nucleus, plasma membrane (interior and exterior), endoplasmic reticulum, and mitochondria **[Extended Data Table 1]**.

Across the target set, T_M,app_ values ranged from 44°C to 64°C, with a mean and median of 49°C and 48°C, respectively. The vast majority of targets melted adequately at temperatures below 60°C, a temperature range over which the NanoLuc reporter is appropriate and cells remain intact. For highly-thermostable proteins such as RAS family GTPases (T_M,app_: 58–63°C) and WDR5 (T_M,app_: 64°C), the StabiLuc reporter was required to obtain an adequate signal and the Extracellular NanoLuc Inhibitor was omitted to prevent inadvertent quenching of the luciferase signal upon thermal lysis. Importantly, BRETSA operated well regardless of target localization, demonstrating that BRETSA Probe 01 equilibrates across intracellular compartments under assay conditions.

To explore the utility of BRETSA for target engagement analysis, a control test ligand was chosen for each target in the panel, and cells were treated with ligand or vehicle to evaluate thermal-protection. However, compared to traditional thermal shift assays, target engagement assessments in BRETSA allowed for an expanded framework for hit-calling analysis. Across the panel, compound engagement perturbed the BRETSA melt in two dimensions; a typical rightward shift in the transition midpoint (an increase in T_M,app_, consistent with classical thermal stabilization) or also a downward shift in the BRET signal at a given temperature (with or without an accompanying shift in T_M,app_). We therefore quantified engagement as the amplitude of BRET inhibition, scored as either (1) the isothermal engagement window (see methods) or (2) the integrated change in BRET across the temperature range, defined as the area between vehicle and compound melt curves (% area-under-the-curve / AUC, see Methods). These qualitative readouts provide a robust binary determination of target engagement across diverse targets and modalities. Across the target-ligand set, the average isothermal engagement window was 2.9-fold, with the largest effects in the 7–8-fold range (PLPro, FKBP12(F36V), and the catalytic domain of USP7, *vide infra*) **[Extended Data Table 1 and Supplementary Data Set 1].** The smallest reproducible windows for confirmed interactions were in the 1.3-fold range (i.e. SOS1, TUBB, and DAPK3). Ninety-eight percent (112/114) of the known unique target-ligand pairs in this study produced isothermal engagement windows ≥ 1.3-fold, which represented a reasonable empirically defined hit-calling threshold **[Extended Data Table 1 and Supplementary Data Set 1].** For most relatively small protein targets (< 50 kDa), the isothermal engagement window was similar regardless of reporter tagging orientation **[Extended Data Table 1 and Supplementary Data Set 1].** To gauge scalability of the assays for HTS settings, we measured Z’ for a representative set of 18 targets spanning a broad range of isothermal engagement windows (1.4 to 4.9-fold) **[Extended Data Fig. 2c–d]**. We found that all targets in the panel with isothermal engagement windows approximately ≥2-fold generated Z’ values ≥0.5, providing a practical threshold for scalability considerations **[Extended Data Fig. 2c–d]**.

When the comprehensive target panel was evaluated via %AUC, interaction calls were in general agreement with the calls from isothermal engagement window. Of 114 target and ligand pairs, 113 (99%) exceeded the statistically determined 10 %AUC threshold (median 40.5 %AUC; range 9.7 to 79.5), and 85% exceeded 20 %AUC **[Extended Data Table 1].** Collectively, BRETSA confirmed modulation of the BRET signal for 113/114 (99%) of the unique interactions, as judged by isothermal engagement window and/or %AUC measurements, with only PARP10 failing both hit-calling metrics.

Though the vast majority of small molecule interactions manifested as stabilizations (e.g. antagonism of the BRET signal), a small subset manifested as destabilizers (e.g. BRET agonists). For example, MELK kinase and P4HB demonstrated clear destabilization when treated with staurosporine or PACMA31, respectively **[Supplementary Data Set 1].** In contrast, ULK3 demonstrated a mixed effect characterized by thermal destabilization with a strong BRET signal reduction at higher temperatures **[Supplementary Data Set 1].**

There was no systemic limitation of BRETSA for large molecular weight proteins. For example, large, multidomain proteins such as PARP14, BRD4, and MET kinase revealed protective compound effects in full-length form, and compound effects caused near-complete BRET inhibition **[Fig. 2e and Supplementary Data Set 1].** However, for some proteins, the protective effect was fractional (e.g. EPHA3, SLK, and POLQ) or barely evident (e.g., USP7 and WRN) **[Fig. 2g and Supplementary Data Set 1].** In these examples, the compound effect was improved when the catalytic domain was segregated for the BRETSA analysis **[Supplementary Data Set 1]**. This domain strategy was leveraged in myriad additional examples such as the kinase domains of TNIK, MERTK and other Ephrin receptors beyond EPHA3, as well as the helicase domain of POLQ **[Fig. 2g and Supplementary Data Set 1]**. For a subset of other targets, the temperature-dependent increase in BRET signal was modest, but the baseline BRET signal was meaningfully elevated even at physiological temperatures (e.g. > 5-fold above the typical BRET signal for no probe controls). In such cases (most strikingly PRMT5, HSP90B1, and PIM3), compound treatment still provided a BRET-reducing effect, supporting direct target engagement **[Supplementary Data Set 1].** In these cases, the BRETSA probe could be pre-associating with a dynamic state of the target, and ligand binding may either change the conformation or cause exclusion of the probe.

Using digitonin-permeabilized cells, BRETSA was also able to resolve engagement for compounds with reportedly poor cellular permeability. Consistent with previous studies^22, 23^, weakly permeable inhibitors of PRMT7 and SETDB1 required digitonin treatment for target engagement in BRETSA **[Extended Data Fig. 2e–f]**.

Because BRETSA detects any perturbation to a target’s thermal stability, its utility should extend beyond small molecules to the detection of protein–protein interactions (PPIs) and higher-order complex formation, which impose their own characteristic shifts on the melt profile. A subset of the validated targets function within protein complexes, prompting us to test how PPIs affect melt behavior and target engagement. In each case, an untagged partner was co-transfected alongside a target tagged with the luciferase **[Fig. 2h].** To explore this concept we selected E3 ligase cereblon (CRBN), kinases BRAF, CRAF, and PI3Kα (p110α, PIK3CA), as well as the transcription factor MYC. Stabilization was evident for a number of selected targets. The DDB1 subunit stabilized CRBN-NanoLuc (∼4 °C), which could be further stabilized by lenalidomide treatment **[Fig. 2i and Extended Data Fig. 2g].** We then evaluated engagement of RAF inhibitors selective for RAS-RAF complexes. KRAS(G12C) destabilized BRAF-NanoLuc (∼3 °C) and compressed the upper asymptote for CRAF-NanoLuc **[Extended Data Fig. 2h and 2i].** Moreover, the RAF-RAS complex ligand LY3009120 provided strong thermal stabilization of both KRAS-RAF complexes **[Extended Data Fig. 2h and 2i]**. To explore the MYC-MAX transcriptional complex, we demonstrated that co-expression of MAX provides strong stabilization of MYC-NanoLuc (>10 °C) **[Fig. 2j and Extended Data Fig. 2j].** Moreover, the p85 subunit (PIK3R1) stabilized the p110α subunit (PIK3CA), observed as both a modest ∼4 °C shift and a strong compression of the upper plateau **[Extended Data Fig. 2k]**. These partner-dependent perturbations both underscore the value of evaluating complex partners during assay development and show that BRETSA can detect protein–protein interactions through ectopic gene dosing experiments.

### Recording longitudinal BRETSA over hundreds of targets en masse

Given the success in adapting BRETSA to a homogeneous format, we next evaluated whether the concept could be scaled more broadly to query target engagement across hundreds of related proteins en masse. Broad spectrum proteomics profiling across entire protein families can identify novel interactions among closely related targets and target families. Proteome-wide thermal proteome profiling by mass spectrometry (TPP/CETSA-MS) has transformed cellular target engagement analysis, enabling largely unbiased target identification across thousands of endogenously expressed proteins^5, 24^. For sufficiently abundant proteins, this capability is unmatched for surveying the engaged proteome. However, for poorly expressed proteins, or proteins recalcitrant to MS analysis, some interactions may remain unresolved. Combined with a focused library of luciferase-tagged genes, BRETSA could complement global proteomics analysis by providing a measurement of thermal stabilization that is independent of endogenous abundance and phase-separation.

To query thermal shift broadly over diverse target proteins with distinct melt profiles, a full range of temperatures would be ideal. Interrogating large target panels therefore represents a challenge, as subjecting individual proteins to a thermal gradient on a well-by-well (endpoint) basis would limit throughput and scale. Unlike most cellular thermal shift assays, the BRETSA readout is non-destructive and compatible with intact cells throughout the entire workflow. The ratiometric feature of BRET also provides a steady signal output, gating the luminescent signal decay that can complicate other tag-based approaches reliant upon raw luminescent intensity^10^. Because denaturation is detected by BRET rather than by separation of soluble and insoluble fractions as an endpoint, the BRETSA measurement does not require cell lysis, centrifugation, or any irreversible processing step. These properties therefore enable longitudinal thermal profiling, in which a single well is read repeatedly as the temperature is incremented stepwise. This single-well analysis of melt behavior would enable a logarithmic increase in scalability for microplate analysis. To evaluate this longitudinal measurement via BRETSA, we compared single-well melt profiles to that of endpoint analysis. Sentinel targets PARP1 and SLK were evaluated in longitudinal vs. endpoint/multi-well mode, and results were comparable (**Extended Data Figure 3**).

We chose the human kinome and the multikinase inhibitor staurosporine to broadly explore single-well melt analysis via longitudinal BRETSA. The kinase library comprised 526 NanoLuc-tagged kinases, including 482 protein kinases and 44 small-molecule kinases, representing highly studied and understudied family members (**Extended Data Table 2**). Constructs were individually reverse-transfected into HEK-293 cells. Each sample was challenged with 10µM staurosporine or DMSO vehicle control by longitudinal BRETSA, canvassing 12 temperatures. Melt midpoints were obtained for 454 of the 526 kinases (86%). Apparent T_M_ ranged from 45.2 to 59.0 °C with a median of 50.9 °C; a further 14 targets melted beyond the applied gradient and are reported as >60 °C **[Fig. 3a; Extended Data Table 2].** Applying a stringent engagement criterion of ≥10% AUC stabilization in both biological replicates, we identified 138 reproducible stabilizing interactions, 129 at protein kinases and 9 at small-molecule kinases **[Fig. 3b–c, Extended Data Table 2].** To orthogonally verify the BRETSA findings, we cross-referenced its staurosporine interactome against two mechanistically independent probe-displacement measurements: intracellular NanoBRET target engagement^25-27^ (300 kinases), and previously reported biochemical competition binding (using purified, recombinant kinases void of ATP)^28^. Of the 94 BRETSA interactions with a matching kinase in the NanoBRET panel, 87 were confirmed (>50% occupancy at 10 µM in NanoBRET). Of the 113 with a matching kinase in the biochemical competition binding panel, 109 were confirmed (Kd < 1 µM in biochemical competition). Among the 235 panel targets addressable by both probe-displacement formats, BRETSA called 90 interactions, of which 89 (99%) were corroborated by at least one orthogonal measurement (EPHA2 being the exception). Notably, a small number of targets were detected by BRETSA and NanoBRET yet scored negative by both the proteomic and biochemical assays, including the lipid kinase PIP4K2C, indicating that cellular BRETSA can detect interactions invisible to purified systems.

**Figure 3.**
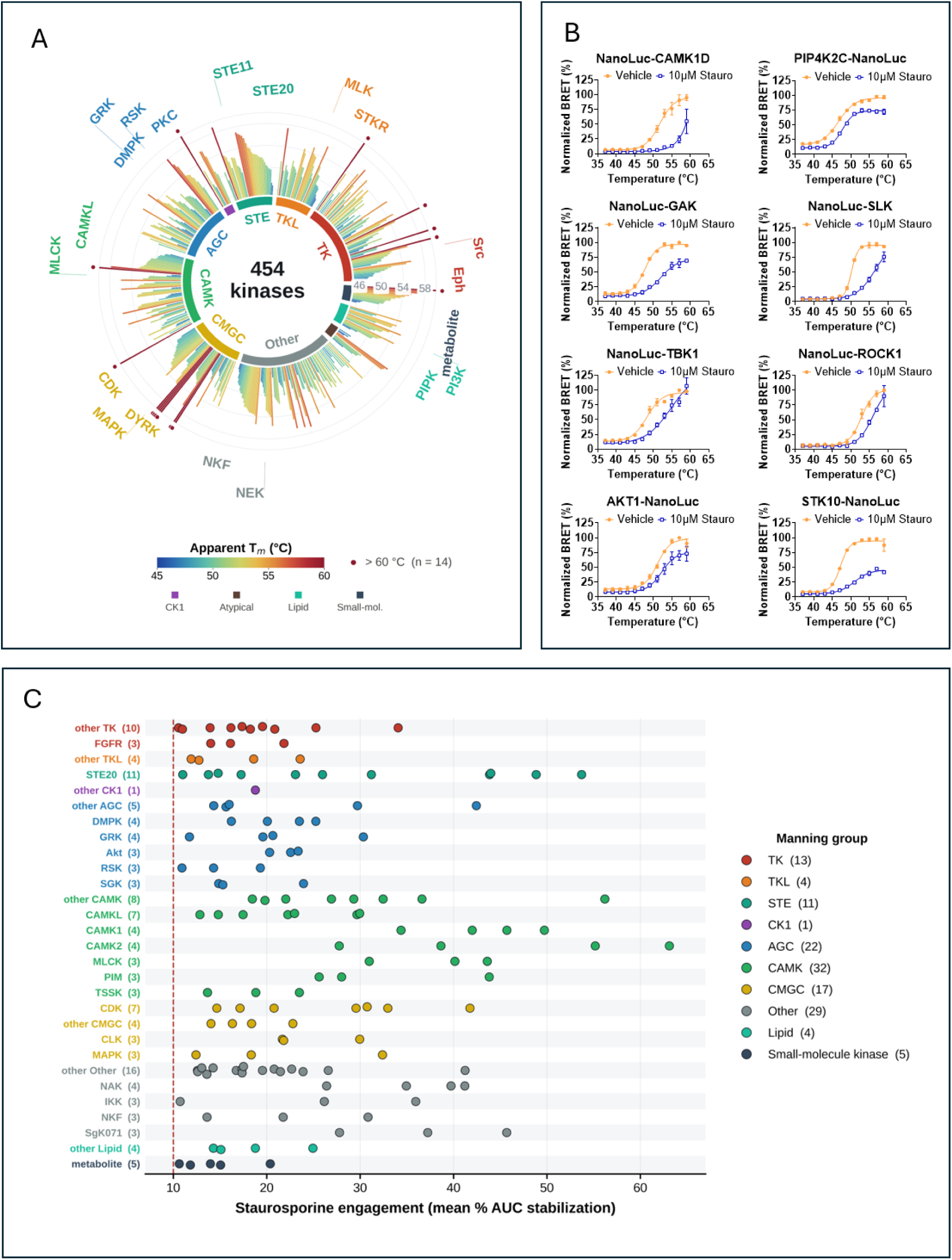
Longitudinal evaluation of the staurosporine interactome. (A) Apparent melting temperatures for 454 NanoLuc-tagged kinases determined by longitudinal BRETSA in the absence of compound, arranged by Manning group and family with targets ordered by ascending T_M_ within each family. Bar length and fill both encode apparent T_M_; the scale is capped at 60°C and the targets exceeding this value are marked with a point outside the ring. (B) Staurosporine interactions detected in BRETSA that were below statistical thresholds for hit-calling in TPP [Savitski 2014]. Single-well melt curves were acquired in two independent biological replicates. For display, BRET values for each target and replicate were linearly scaled between the plate-specific NanoLuc control signal (unfused NanoLuc with Transfection Carrier DNA) at 37°C and the maximum vehicle-treated signal across the temperature gradient, with the same floor and ceiling applied to the vehicle and staurosporine curves. This scaling preserves the shape of both curves and the relative area between them, up to a single scaling constant for each target and replicate, while correcting for differences in absolute BRET signal between experiments. %AUC stabilization **[Extended Data Table 2]** was calculated from raw BRET values prior to scaling and is not recalculated from the plotted curves. Points show the mean of the two replicates and error bars show the range. (C) Staurosporine engagement for the 138 targets meeting the hit criterion of ≥10% AUC stabilization in both biological replicates, grouped by Manning kinase family and colored by group. Values are the mean %AUC of the two biological replicates; the dashed line marks the 10% threshold.

Finally, we compared this interaction landscape with the reported proteome-wide TPP by mass spectrometry using K562 cell lysates^5^. Within the kinase family, BRETSA and TPP identified 23 common staurosporine stabilizing interactions, while each method identified a similar number of interactions undetected by the other **[Extended Data Table 2].** Some such discrepancies could be explained by the use of K562 cell extracts and dilution of cellular co-substrate (ATP) in TPP, whereas BRETSA analysis was performed in live cells, where co-substrate concentration may vastly exceed ATP K_m_ for some kinases. Of the 138 staurosporine interactions identified by BRETSA, 115 were not reported by TPP. For 26 of these, TPP measured the kinase in both replicates but detected no stabilization. The remainder (89) fell outside the detection limits of the proteomic experiment, having either never been identified in the K562 proteome or yielding no analyzable melting curve, reflecting the limits of protein abundance and curve-fit significance at proteome scale. It is noteworthy that a number of the BRETSA-specific staurosporine interactions fell within the lesser studied / dark fraction of the kinome^29^, including the TSSK family, the STK32 subfamily, and multiple CAMK and DCLK members, targets likely unrepresented in K562 lysates. In chemoproteomics studies, TPP could be leveraged for unbiased identification of target families and related pathways, while BRETSA could be leveraged for more comprehensive dissection of engagement at lesser studied / low abundance proteins within those families and pathways.

### Isothermal BRETSA provides accurate potency stratification

The binary BRETSA analyses described above establish target engagement, but with limited quantitation. To rank-order affinity and stratify analog series for medicinal chemistry, engagement is ideally resolved as a function of compound concentration. Isothermal dose-response analysis addresses this by fixing the temperature and varying compound concentration, producing sigmoidal curves from which IC_50_ values can be extracted. However, because ligand binding is thermodynamically coupled to protein folding, we reasoned that the potency measured in a thermal shift assay should depend on the temperature at which it is assessed. BRETSA matrix data (in which compound dose and temperature are varied simultaneously) were re-evaluated, resolving potency as a continuous function of thermal challenge **[Supplementary Data Set 2]**. Isothermal IC_50_ values were temperature-dependent for a wide array of reversible inhibitors **[Supplementary Data Set 2],** weakening systematically as assay temperature was raised. We observed this across the majority of reversible interactions in our panel, including more striking examples such as MAPK14/AMG548, p53(Y220C)/rezatapopt, WDR5/OICR-9429, and SHP2/RMC-4550 **[Fig. 4a–c and Supplementary Data Set 2]**.

**Figure 4.**
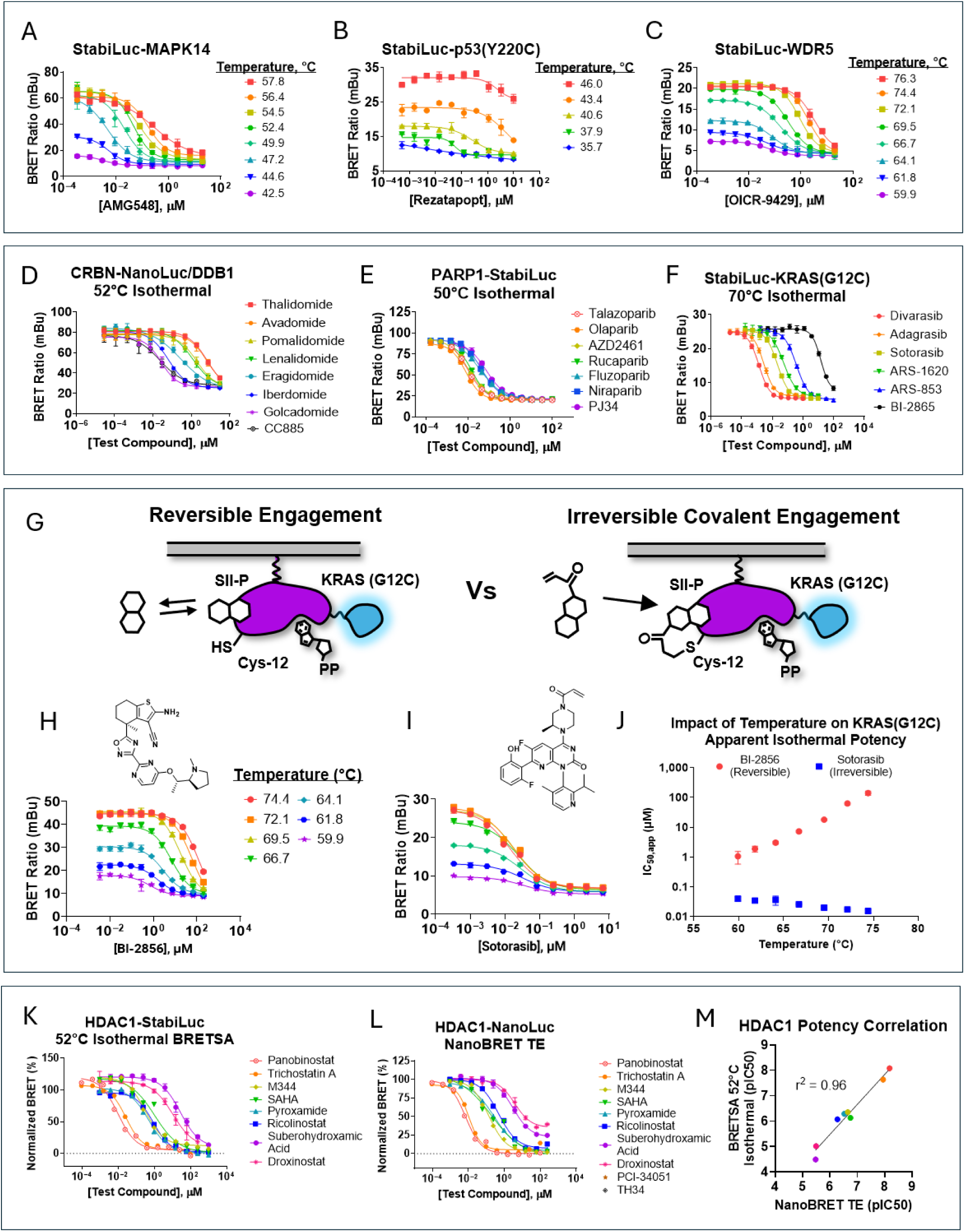
Isothermal BRETSA for potency determination. (A-C) Example temperature dependent isothermal engagement potencies for MAPK14 (A), p53(Y220C) (B), and WDR5 (C). Data are representative of experiments performed twice where individual data points are the mean (± range) of 2 technical replicates. (D–F) Isothermal compound engagement profiles for CRBN (D), PARP1 (E), and KRAS(G12C) (F). Data are representative of experiments performed 2 times (CRBN and KRAS(G12C)) or 3 times (PARP1), where individual data points are the mean (± SD) of 4 technical replicates. (G) Schematic comparison of reversible vs irreversible covalent engagement of the KRAS(G12C) Switch-II pocket (SII-P). (H-J) Temperature dependent isothermal potency for KRAS(G12C) SII-pocket ligands. BRETSA Isothermal potencies for reversible ligand BI-2865 (H) and irreversible SII-pocket ligand sotorasib (I). Data are representative of experiments performed twice, where individual data points represent the mean (± range) of technical duplicates. (J) Temperature dependent isothermal potencies for ligands from panels H and I. Data points are the mean (± range) from two independent experiments. (K–M) HDAC1 ligand potency profiles for isothermal BRETSA (K) vs NanoBRET TE (L). Data are representative of experiments performed twice, where individual data points represent the mean (± range) of 2 technical replicates. (M) Correlation plot of potency (pIC50) values between isothermal BRETSA and NanoBRET TE. Individual data points are the mean of experiments performed twice.

At a fixed temperature near the target’s T_M,app_, BRETSA consistently produced sigmoidal dose-response curves suitable for compound potency ranking **[Supplementary Data Set 2],** providing further evidence of bona fide engagement. Additional example target protein/ligand pairs were selected to evaluate the detection sensitivity limits for apparent isothermal potency. Targets where compound profiles were established include epigenetic targets PARP1 and HDAC1, the E3 ligase CRBN, kinases MAPK14, AURKB, and ABL1, the ATPase/chaperone HSP90AB1, the catalytic domain of USP7, and the GTPase KRAS(G12C) **[Fig. 4d–f, 4k and Extended Data Fig. 4]**. BRETSA resolved potencies spanning greater than 5 orders of magnitude, exemplified by USP7(207-650) with USP7-IN-11 (130pM) **[Extended Data Fig 4e]** and HDAC1 with suberohydroxamic acid (33µM) **[Fig. 4k and Extended Data Fig 4j]**. Isothermal analysis enabled potency determination for not only orthosteric inhibitors but also allosteric inhibitors for a number of proteins across at least 5 classes (e.g., Abl1, kinase; USP7, deubiquitinase; p53(Y220C), transcription factor; SHP2, phosphatase; WRN, helicase) **[Fig. 4b, Extended Data Fig. 4c and 4e, and Supplementary Data Set 2]**.

We next explored the behavior of covalent target engagement in BRETSA. Unlike reversible engagement, irreversible covalent inhibition is governed by the kinetics of adduct formation, in contrast to binding equilibrium. Accordingly, irreversible covalent interactions should be insulated from perturbation by thermal challenge. To query this relationship, irreversible covalent ligands and reversible ligands were compared against the same target protein in BRETSA **[Fig. 4g]**. The switch-II pocket inhibitors sotorasib and BI-2865 were tested against the G12C mutant of KRAS **[Fig. 4h–j and Extended Data Fig. 4g]**. Sotorasib did not show temperature-dependent effects on drug IC50 values over a broad temperature range, consistent with irreversible occupancy. In contrast, the reversible switch-II pocket inhibitor BI-2865 demonstrated potencies that were highly temperature sensitive. Target engagement with sotorasib was extended to G12C alleles of other RAS family members (NRAS, HRAS), supporting switch-II pocket vulnerability **[Extended Data Fig. 4h]**. We extended a similar mechanistic study to Aurora kinase A, where covalent and reversible ATP competitive inhibitors could be resolved **[Extended Data Fig. 4i]**. Cumulatively, these data support the hypothesis that the isothermal potency of irreversible covalent ligands will be relatively insensitive to temperature, likely due to the fact that the covalent bond is established prior to thermal challenge.

To evaluate rank-order target engagement accuracy, we compared isothermal BRETSA IC50 values to potencies from NanoBRET target engagement (TE) tracer assays—an orthogonal method that directly measures compound occupancy via competitive displacement in cells^17, 25, 27, 30^. Across compound panels for HDAC1 (8 compounds), PARP1 (7 compounds), and CRBN (8 compounds), BRETSA and NanoBRET TE IC_50_ values were highly correlated (R² = 0.96, 0.97, and 0.95, for HDAC1, PARP1, and CRBN, respectively) **[Fig. 4k–m and Extended Data Fig. 4j–l]**. Potency offsets between NanoBRET TE and BRETSA isothermal potencies varied by target and compound and ranged from no difference (olaparib for PARP1) to 11-fold (Pomalidomide for CRBN), generally suggesting that BRETSA isothermal potencies have a tendency to be marginally weaker after thermal challenge compared to NanoBRET TE potencies at physiological temperatures **[Fig. 4k–m and Extended Data Fig. 4j–l].** Despite the potency offsets, these comparisons confirm that isothermal BRETSA faithfully reports relative target occupancy and support the utility of BRETSA to stratify an SAR series through potency analysis. To address whether any of the potency offsets between these formats were caused by ectopic overexpression of the tagged target, we confirmed that transient transfection and endogenous-level (CRISPR knock-in) expression of NanoLuc fusions produced equivalent BRETSA pharmacology for MAPK14, BRD4, CDK2, and CDK4, with isothermal potency values in close agreement across expression systems **[Extended Data Fig. 4m–p]**. These results support that target concentrations in the transiently expressed BRETSA assays are sufficiently low and are not causing potency offsets in binary target engagement studies.

### BRETSA is more sensitive than DSF and luciferase-based methods coupled to aggregation

Apparent potencies derived from thermal shift are routinely weaker than those obtained by direct binding methods, limiting the dynamic range over which engagement can be measured and rank-ordered^31^. Having established a sensitive isothermal potency analysis in BRETSA, we sought to compare these potencies to those generated in other biophysical methods for biochemical or cellular target engagement **[Fig 5a]**. To benchmark isothermal BRETSA against the most established and widely used acellular biophysical thermal shift method, we compared it to DSF, which shares BRETSA’s direct denaturation-detection principle but requires micromolar purified protein **[Fig. 5a]**. Across four matched targets (MAPK14, PARP1, AURKB, and ABL1), DSF isothermal potencies were consistently >1 µM and compressed near the assay protein concentration (micromolar) **[Fig. 5c–d, Extended Data Fig. 5a–d]**. In contrast, BRETSA resolved a far wider range for each protein in the study **[Fig. 4e**, **Fig. 5b; Extended Data Fig. 4a–c and 4k]**: for MAPK14 BRETSA discriminated a >250-fold span (doramapimod, 3nM; SB239063, 800nM, **Fig. 5b and Extended Data Fig. 4a**) that DSF compressed into a 3–5-fold range **[Fig. 5c and Extended Data Fig. 5a]**, and for ABL1 it detected ponatinib at 2nM **[Extended Data Fig. 4c]** versus 3.5µM by DSF (>1,000-fold) **[Extended Data Fig. 5d]**.

**Figure 5.**
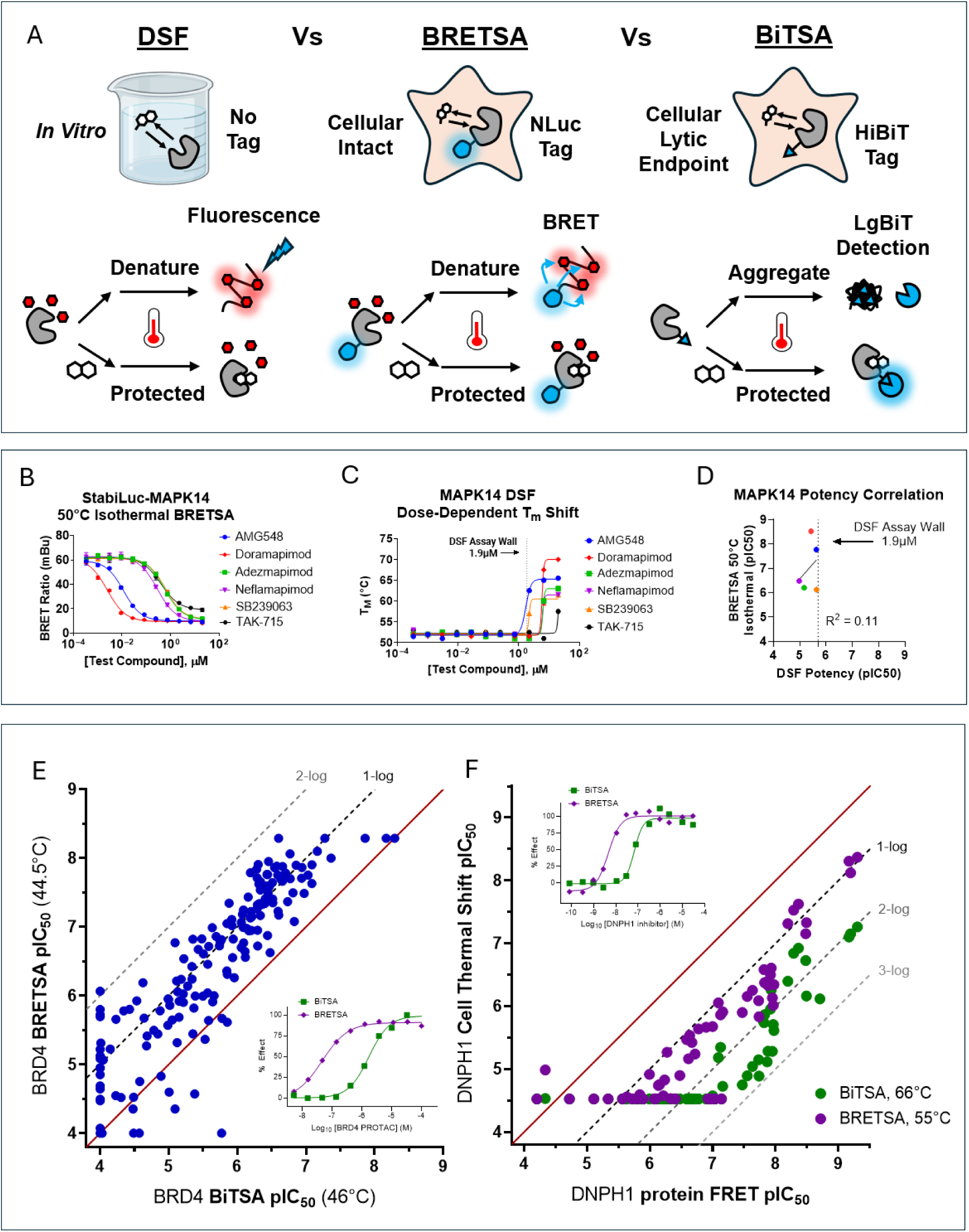
BRETSA compared to other thermal shift methods. (A) Schematic overview comparing BRETSA to other thermal shift assays such as DSF and BiTSA. (B) MAPK14 BRETSA isothermal potency profile for a series of kinase ligands. Data are representative of experiments performed twice where individual data points are the mean (± SD) of 4 technical replicates. (C) MAPK14 DSF potency profile using dose-dependent T_M_ shifts for the same ligand series as in panel B. Data are representative of experiments performed twice, where T_M_ values at each dose were extracted from melt curves collected in technical singlicate. (D) Correlation plot of potency (pIC50) values between isothermal BRETSA and DSF. Individual data points are the mean of experiments performed twice. (E) Correlation plot of potency (pIC50) values between the lowest robust screening temperature (RZ’ >0.5, **Extended Data Fig. 6b**) of isothermal BRD4 BiTSA (46°C) and isothermal BRD4 BRETSA (44.5°C) for 193 compounds and PROTACs containing BET-domain pharmacophores. Each IC50 was calculated from a 9- or 10-point dose response n=1 normalized to on-plate controls, with representative BRD4 PROTAC response shown inset. (F) Correlation plot of potency (pIC50) values between recombinant DNPH1 protein probe displacement FRET binding assay and indicated isothermal Cell Thermal Shift (DNPH1 BiTSA at 66°C green and DNPH1 BRETSA at 55°C purple) for 60 DNPH1 inhibitors spanning a range of potencies. Each IC50 was calculated from a 12-point dose response n=1 normalized to on-plate controls, with representative DNPH1 inhibitor response shown inset.

BRETSA and DSF potencies consistently demonstrated poor correlations **[Fig. 5d and Extended Data Fig. 5b–d]**, most likely reflecting the assay wall imposed by DSF due to the use of micromolar protein concentrations. Despite poor correlations in apparent test compound potency, the apparent melting temperature observed in BRETSA was either similar or marginally lower than in DSF for the 3 proteins where the full length ORF was used in both methods (MAPK14, PARP1, and AURKB) **[Extended Data Fig. 5e]**, demonstrating that the NanoLuc or StabiLuc tags used in BRETSA do not impart a stabilizing effect on the fused protein.

Owing to simplicity and scalability in assay design, luciferase-based thermal shift methods have become increasingly standardized in cell-based workflows^7, 9, 10, 32^. However, each has a fundamental dependency on aggregation. Because BRETSA does not require aggregation, we asked how its behavior differs from aggregation-based methods. Using the MAPK14 model system, BRETSA potencies were nearly 10-fold stronger than those reported using direct NanoLuc luminescence (NaLTSA)^7^. Next, we systematically compared BRETSA to aggregation-based cell thermal shift wherein the thermostable fraction of target protein is assessed via a split-reporter HiBiT tag (BiTSA). Robust and scalable BiTSA assays benefitted from the physical separation of the heated sample into a separate detection plate^32^. We compared the behavior of StabiLuc-tagged BRD4 in BRETSA with HiBiT-tagged BRD4 in BiTSA. Despite similar BRD4 melting profiles, a robust isothermal BiTSA assay required >80% target unfolding (≥46°C), while BRD4 BRETSA enabled a robust screening format at lowered temperatures (≥44.5°C) **[Extended Data Fig. 6a–b]**. Configuring such assays at their lowest robust screening temperatures, we determined IC50 for 193 compounds containing BET-domain pharmacophores spanning both inhibitors and PROTACs and observed a roughly 10-fold improvement in apparent potency in BRETSA compared to BiTSA **[Fig. 5e and Extended Data Fig. 6c]** driven by lowered assay temperature **[Extended Data Fig. 6d–e]**. We next extended the evaluation to a recently reported series of nucleoside hydrolase DNPH1 inhibitors^33^.

Quantification of thermostable HiBiT-tagged DNPH1 using a BiTSA assay demonstrated a DNPH1 T_agg_ of 63.5 °C and required an isothermal temperature of at least 66 °C to support robust BiTSA compound screening, wherein DNPH1 inhibitors were observed to be 100-to- 1000-fold weaker than reported by a recombinant protein probe-displacement FRET assay **[Extended Data Fig. 7a and 7c]**. DNPH1-StabiLuc BRETSA revealed a markedly lower T_m,app_ of 47.7°C, facilitating robust BRETSA compound profiling at temperatures as low as 55°C **[Extended Data Fig. 7b]**. BRETSA screening of 60 DNPH1 inhibitors at 55 °C demonstrated ∼10-fold (1-log) enhancement of apparent potency compared to BiTSA **[Fig. 5f and Extended Data Fig. 7d]**, resolving DNPH1 ligand potency closer to IC50 determined in a purified-protein setting and quantifying target engagement of weaker DNPH1 ligands which could not be resolved by aggregation-based BiTSA. For the targets evaluated here, BRETSA generally resolved target engagement potency across a wider range than DSF or aggregation-based methods.

### Measuring cooperativity at induced ternary complexes in cells

Having observed the ability to query PPIs in BRETSA, we evaluated how PPIs could influence drug potency at higher-order complexes. A growing class of proximity-based modalities function by stabilizing ternary protein–drug–protein complexes, including both molecular glues and heterobifunctional modalities such as RIPTACs^34^ **[Fig. 6a]** and TCIPs^35^. The thermodynamic cooperativity (α) of ternary complex formation is a critical determinant of mechanism, selectivity, and therapeutic index. While apparent α (α_App_) is commonly assessed using pure proteins or domains, we reasoned that BRETSA could quantify α_App_ within the cellular milieu by comparing the isothermal IC50 of a compound at a StabiLuc-tagged protein in the presence versus absence of an overexpressed partner protein: α_App_ = IC50(binary) / IC50(ternary).

**Figure 6.**
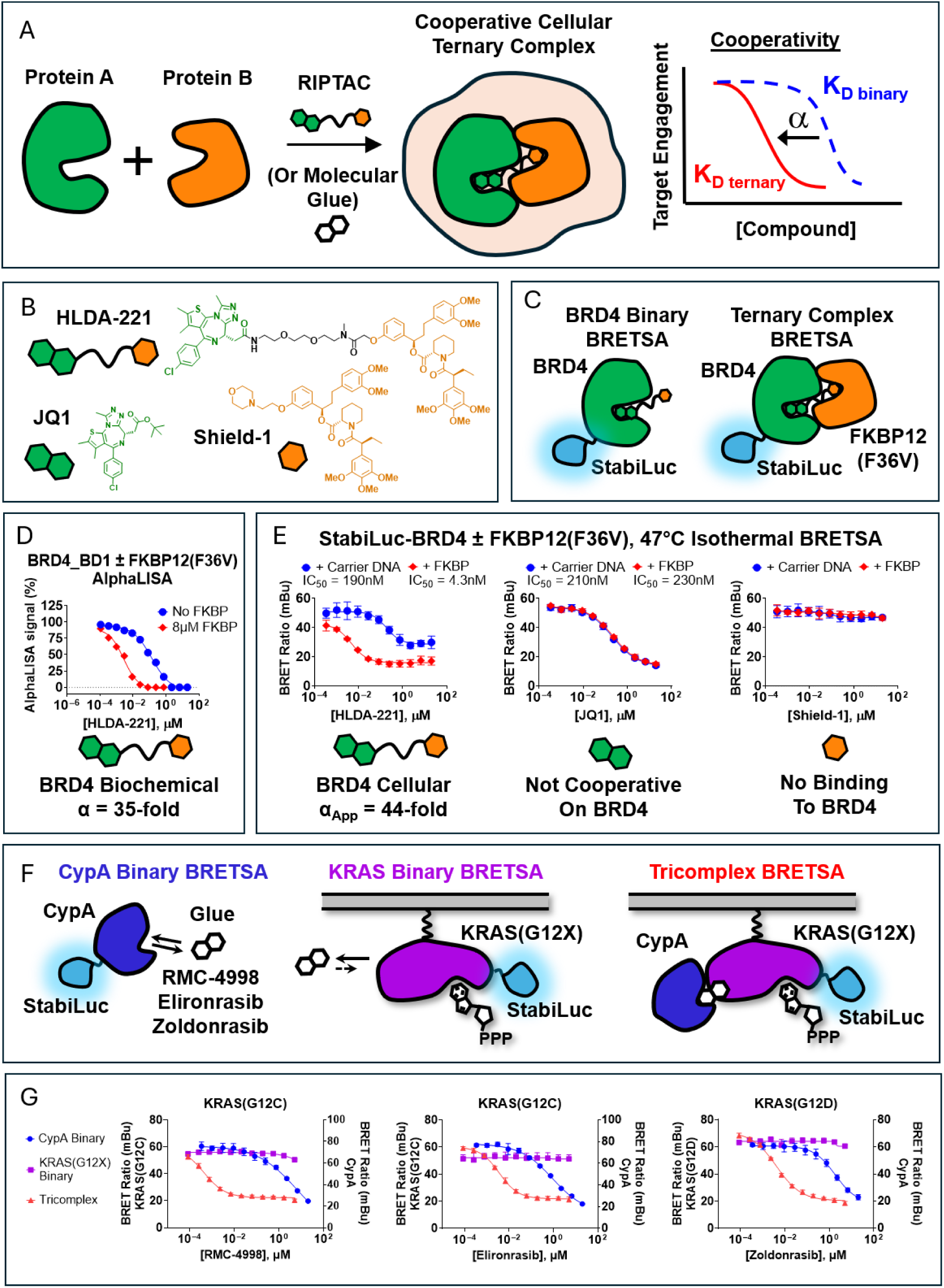
Cellular TE and cooperativity for a model RIPTAC and non-degrading molecular glues. (A) Schematic depiction of the RIPTAC or molecular glue mechanism of action. (B) Chemical structures of the HLDA-221 BRD4/FKBP12(F36V) RIPTAC, as well as the monovalent control compounds JQ1 and Shield-1 used in the study. (C) Isothermal BRETSA assay schematics for the determination of HLDA-221 cellular cooperativity for the BRD4/FKBP12(F36V) ternary complex. (D) Biochemical cooperativity data for HLDA-221 ternary complex with BRD4_BD1 and FKBP12(F36V) replotted from *Raina et al. 2024*. (E) StabiLuc-BRD4 isothermal BRETSA (47°C) in the presence and absence of FKBP12(F36V) co-expression for the compounds depicted in panel B. Data are representative from experiments performed twice. Individual data points represent the mean (± SD) of 4 technical replicates. Apparent cooperativity (α_App_) was calculated from the ratio of the binary to ternary IC50 values. (F) Isothermal BRETSA assay schematics for the determination of cellular cooperativity for the RAS-ON CypA-dependent molecular glues RMC-4998, Elironrasib, and Zoldonrasib in CypA deficient HAP1 cells. Isothermal BRETSA for CypA-StabiLuc, StabiLuc-KRAS(G12C), and StabiLuc-KRAS(G12D) were performed at 55°C, 71°C, and 71°C, respectively. (G) Isothermal BRETSA potencies for RMC-4998, Elironrasib, and Zoldonrasib using the assay configurations depicted in panel F. Data are representative of experiments performed twice, where individual data points are the mean (± SD) of 4 technical replicates).

Early proof-of-concept studies on the RIPTAC mechanism utilized HLDA-221, which leverages BRD4 as the essential protein and an engineered FKBP12(F36V) mutant as the target protein. Structurally, HLDA-221 comprises BRD4 and FKBP12(F36V) targeting moieties **[Fig. 6b]** and previously demonstrated cooperative binding *in vitro^34^* **[Fig. 6d]**. After confirming binary interactions with BRD4 and FKBP12(F36V) using monovalent controls **[Fig. 6e and Extended Data Fig. 8e]**, we evaluated the influence of FKBP12(F36V) overexpression on melt behavior and target engagement at BRD4 **[Fig. 6c, 6e, and Extended Data Fig. 8a–d]**. Relative to isolated StabiLuc-BRD4, HLDA-221 demonstrated significantly enhanced isothermal potency at BRD4 when FKBP12(F36V) was overexpressed (IC50 4 nM) **[Fig. 6e, Extended Data Fig. 8b]**, consistent with cooperative ternary complex formation, whereas the potency of monovalent JQ1 was unchanged **[Fig. 6e, Extended Data Fig. 8c]**. Comparing the ternary and binary potencies of HLDA-221 at BRD4 yielded an α_App_ of ∼44-fold **[Fig. 6e, Extended Data Fig. 8b]**, in excellent agreement with the biochemical cooperativity reported previously^34^. Moreover, pre-treatment of StabiLuc-BRD4/FKBP12(F36V) expressing cells with excess Shield 1 ligand prior to treatment with HLDA-221 caused reversion of the apparent isothermal potency back to that observed in the absence of FKBP12(F36V) overexpression **[Extended Data Fig. 8f]**, providing further support for the cooperative mechanism of action.

We next explored KRAS(ON) inhibitors that exploit a distinct tricomplex mechanism, recruiting the endogenous chaperone cyclophilin A (CypA) to form a KRAS–compound–CypA ternary complex that blocks effector interactions^36-38^. RMC-4998 and elironrasib target KRAS(G12C), while zoldonrasib (RMC-9805) targets KRAS(G12D). We applied BRETSAs to measure the binary interactions of these glues with CypA and KRAS mutants, as well as the potency of CypA-dependent KRAS/glue complex formation **[Fig. 6f and Extended Data Fig. 8g]**. To assess these interactions, a CypA(−) knockout cell line (Hap1) was utilized. RMC-4998 and elironrasib showed no detectable binary engagement at StabiLuc-KRAS(G12C) in CypA(−) Hap1 cells, consistent with the low intrinsic affinity of these compounds for KRAS in the absence of CypA **[Fig. 6g and Extended Data Fig. 8h]**. Binary engagement in cells expressing CypA-StabiLuc (55 °C) yielded IC50 values of 6.2µM and 1.1µM, respectively, consistent with the weak to modest affinities of these ligands for the chaperone^36, 37^ **[Fig. 6g and Extended Data Fig. 8h]**. However, when CypA was co-expressed with StabiLuc-KRAS(G12C), both compounds produced significant dose-responsive stabilization of KRAS(G12C), with isothermal potencies of 0.4 nM and 0.6 nM, respectively **[Fig. 6g and Extended Data Fig. 8h]**. The cooperativity of these interactions exceeded 1,000-fold, regardless of whether the CypA or KRAS interaction was used as the binary reference. We observed similarly extreme cooperativity for zoldonrasib, where BRETSA on KRAS(G12D) with CypA co-expression produced a markedly enhanced dose-response relative to CypA alone **[Fig. 6g and Extended Data Fig. 8h]**, again consistent with α_App_ >1,000. Cumulatively, these values across both the RIPTAC and molecular glue systems demonstrate that BRETSA resolves cooperativity across a wide dynamic range and provides a cellular platform to interrogate biomolecular cooperativity without purified proteins.

### Limitations of BRETSA

BRETSA is not without limitations. The method requires a genetically encoded reporter, a constraint shared with other reporter-based platforms. This limits the utility of BRETSA to cell systems compatible with recombinant engineering. Similar to other thermal shift assays, a fraction of ligand interactions are undetectable, and application to non-globular proteins is generally problematic. Integration of thermal perturbations in high-throughput settings can be challenging, requiring specialized instrumentation and workflows in library screening applications.

### Summary

Here we report the methodological framework for BRETSA and demonstrate broad applications in target engagement analysis. By exploiting a non-destructive, denaturation-based mechanism, BRETSA offers improved scalability via longitudinal analysis and extends sensitivity compared to other thermal shift methods. BRETSA has been successfully applied to target engagement for a wide array of chemical modalities. Extending on traditional binary pharmacology studies, BRETSA is ideally suited for assessments of drug cooperativity at biomolecular complexes without pure proteins or advanced target pre-knowledge. Future directions for BRETSA are broad but should entail extensions into non-globular proteins and hit finding for intractable drug targets.

## Methods

### DATA AVAILABILITY

The authors declare that the main data supporting the findings of this study are available within the article and its Supplementary Information files. Requests for further information, resources, and reagents should be directed to and will be fulfilled by the lead contact Matthew B. Robers.

### EXPERIMENTAL MODELS AND SUBJECT DETAILS

HEK-293 cells (ATCC) were cultured in DMEM (Gibco) + 10% fetal bovine serum (FBS) (Seradigm). K562 cells (ATCC) were cultured in RPMI (Gibco) + 10% FBS. Parental HAP1 cells (Horizon Discovery) as well as HAP1 CypA(–) cells (Horizon Discovery) were grown in Iscove’s Modified Dulbecco’s Medium (IMDM). All cells were incubated in a humidified, 37°C/5% CO_2_ incubator. Three CRISPR edited cell lines were obtained from Promega including MAPK14-NanoLuc (HEK-293, Cat.# CS302303), NanoLuc-BRD4 (HEK-293, Cat.# CS302313), CDK2-NanoLuc (HEK-293, Cat.# CS302301), or NanoLuc-CDK4 (K562, Cat.# CS302323). HEK-293 cells endogenously expressing HiBiT-BRD4 were prepared previously^32^. SUM149PT cells endogenously expressing HiBiT-DNPH1 were prepared previously^33^ using CRISPR/Cas9 editing and grown in Nutrient F12 Ham (Sigma N6658) medium supplemented with 1% FBS.

### METHOD DETAILS

#### Denaturation Sensitive Probes, Chemical Inhibitors, and Other Chemical Reagents

Sypro Orange (5000X) and Sypro Red (5000X) were from ThermoFisher Scientific. BRETSA™ Probe-01 (1000X) was from Promega Corporation. All chemical inhibitors were from common vendors such as Selleckchem, MedChemExpress, or Tocris. Concentrated stock solutions of chemical inhibitors were prepared in DMSO at 100X–1000X and then either diluted in Opti-MEM without phenol red (Gibco) or dispensed directly from DMSO (as noted).

#### Cell Transfections

All luciferase fusion proteins in this study were encoded in pFN31K (N-terminal NanoLuc^®^), pFC32K (C-terminal NanoLuc^®^), pSL53K (N-terminal StabiLuc™), or pSL54K (C-terminal StabiLuc™) expression vectors. All fusion proteins were encoded with flexible GSSG linkers between the target protein ORF and the luciferase ORF with the exception of HDAC1, which used an alternate linker (SWTWEGNKWTWK). All unfused proteins were encoded in pFN5K expression vectors. Target protein ORFs generally corresponded to Uniprot isoform 1 (or named mutants thereof), unless otherwise noted in Extended Data Tables 1 and/or 2. All transfections used either FuGENE 4K (Promega) or FuGENE HD (Promega) interchangeably according to the manufacturer’s protocol. Unless otherwise noted, target protein/luciferase fusion constructs were diluted into Transfection carrier DNA (Promega) or an expression vector for an unfused partner/regulatory protein at a mass ratio of 1:9 (mass/mass) in Opti-MEM without phenol red (Gibco), after which FuGENE reagent was added at a ratio of 1:3 (µg DNA: µL FuGENE reagent). For example, for a 1mL size transfection complex, 1µg each of the Target protein/luciferase fusion DNA was combined with 9µg Transfection Carrier DNA in 1mL of Opti-MEM. 1 part (vol) of FuGENE HD complexes thus formed were combined with 20 parts (vol) of HEK-293 cells suspended at a density of 2 x 10^5^ per mL in DMEM + 10% FBS (Seradigm). Cell lines beyond HEK-293 were grown and transfected in alternative cell culture media as described above. Cell/transfection complex suspensions were grown in a humidified 37°C/5% CO_2_ incubator for 18−24 hr unless otherwise noted.

#### BRETSA Protocol for 384-well format

HEK-293 cells (unless otherwise noted) transfected as described above were washed with 1X DPBS (Gibco) and dissociated from the flask by trypsinization with TrypLE Express (GIBCO) or 0.05–0.25% Trypsin (Gibco). Trypsinized cells were harvested by quench with an equal volume of FBS-containing cell culture medium, after which the suspension was pelleted and the supernatant removed. The pelleted cells were resuspended in Opti-MEM without phenol red (Gibco) at a cell density of 500,000 cells/mL. Test compound was dispensed directly from a concentrated DMSO stock solution to Bio-Rad Hard-Shell 384-well PCR plates (Cat.# HSP3805, white-well) using either a droplet digital dispenser (Tecan D300e) or an acoustic dispenser (Beckman Echo 650). Test compounds were dispensed either at fixed doses or as dilution series (as noted). BRETSA™ Probe-01 and cell suspension were dispensed to the plates using two different methods interchangeably depending upon the workflow (methods 1 and 2, respectively). For method 1, BRETSA™ Probe-01 was dispensed directly from the 1000X stock solution to the PCR plates containing test compound using either a droplet digital dispenser (Tecan D300e) or an acoustic dispenser (Beckman Echo 650), after which 20µL of cell suspension (prepared above) was seeded per well using a bulk reagent dispense (Integra WellJet or Agilent BioTek MultiFlo FX). For method 2, BRETSA™ Probe-01 was added directly to the cell suspension (prepared above) to a final concentration of 1X, after which 20µL of cell/probe suspension was seeded per well using a multichannel repeater pipette. When tested, Sypro dyes were dispensed directly from the 5000X stock using method 1 using a Tecan D300e. PCR plates containing cells, test compound, and BRETSA™ Probe were capped with a plastic lid (Nunc Microplate lid, ThermoFisher Scientific) and incubated for 1 hr (unless otherwise noted) in a humidified 37°C/5% CO_2_ incubator. Following equilibration, the plates were removed from the incubator, the lid was removed and the plate was sealed with an aluminum adhesive plate seal. Sealed plates were subjected to thermal challenge using a gradient thermal cycler (Bio-Rad PTC Tempo, 384-well format). Thermal challenges were executed for 3 min including a pre-heated lid at 5°C higher than the highest temperature in the challenge. Unless otherwise noted, thermal challenges were either gradients from 37°C–60°C (live-cell range), or a single temperature (isothermal, chosen as a balance between isothermal engagement window and sensitivity to potency and often near the EC_80_ of the melt curve). After thermal challenge, the thermocycler protocol was used to rapidly return the plate to 21°C after which TarSeer™ Detection Substrate (Promega) and TarSeer™ Debris Quencher (Promega) were added according to the manufacturer’s recommended protocol. Assay plates were placed into the 384-well plate adapter for cross-talk reduction (Promega Cat.# GM3055), and filtered luminescence was measured on a GloMax^®^ Discover luminometer equipped with 450nm BP filter (donor) and 600nm LP filter (acceptor), using 0.5 s integration time. Raw BRET ratios were calculated by dividing the acceptor counts by the donor counts. Milli-BRET units (mBU) were calculated by multiplying the raw BRET values by 1000. When normalized BRET was used (for example to compare compound isothermal potencies), mBRET values were normalized using Equation 1;

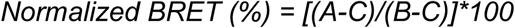

Where A = mBRET in the presence of test compound, B = mBRET in the presence of vehicle, and C = mBRET in the presence of a saturating dose of the test compound to define the bottom. Apparent melt temperature (T_M_) values were determined by plotting the mBRET values as a function of temperature and fitting the Boltzmann sigmoidal equation available in GraphPad Prism 11 (Equation 2);

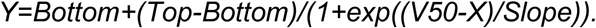

Apparent isothermal potencies (IC_50_) were determined by plotting the mBRET values or normalized BRET values measured at a single temperature (as noted) as a function of compound dose and then fitting the sigmoidal dose-response (variable slope) equation available in GraphPad Prism 11 (Equation 3);

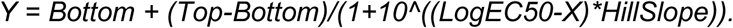

#### BRETSA Protocol for 96-well format

HEK-293 cells (unless otherwise noted) transfected as described above were washed with 1X DPBS (Gibco) and dissociated from the flask by trypsinization with TrypLE Express (GIBCO) or 0.05–0.25% Trypsin (Gibco). Trypsinized cells were harvested by quench with an equal volume of FBS-containing cell culture medium, after which the suspension was pelleted and the supernatant removed. The pelleted cells were resuspended in Opti-MEM without phenol red (Gibco) at a cell density of 400,000–450,000 cells/mL depending on the workflow (see below). Test compound was dispensed to Bio-Rad Hard-Shell 96-well PCR plates (Cat.# HSP9655, white-well) either directly from DMSO using a droplet digital dispenser (Tecan D300e) or from a 10X intermediate stock prepared in Opti-MEM without phenol red. Test compounds were dispensed either at fixed doses or as dilution series (as noted). BRETSA™ Probe-01 and cell suspension were dispensed to the plates targeting a 1X concentration (unless otherwise noted) using 4 different methods interchangeably depending upon the workflow (methods 1–4, respectively). For method 1, test compound was dispensed directly from DMSO and BRETSA™ Probe-01 was dispensed directly from the 1000X stock solution to the PCR plates containing test compound using a droplet digital dispenser (Tecan D300e), after which 100µL of cell suspension (prepared above, 400,000 cells/mL) was seeded per well using a multichannel repeater pipette. For method 2, test compound was dispensed directly from DMSO and BRETSA™ Probe-01 was added directly to the cell suspension (prepared above, 400,000 cells/mL) to a final concentration of 1X, after which 100µL of cell/probe suspension was seeded per well using a multichannel repeater pipette. For method 3, 10µL of a 10X intermediate stock test compound was dispensed using a multichannel repeater pipette and BRETSA™ Probe-01 was added directly to the cell suspension (prepared above, 450,000 cells/mL) to a final concentration of 1.11X, after which 90µL of cell/probe suspension was seeded per well using a multichannel repeater pipette. For method 4, 10µL of a 10X intermediate stock test compound was dispensed using a multichannel repeater pipette and BRETSA™ Probe 01 was dispensed directly from DMSO on top of the test compound, after which 100µL of cell suspension (prepared above, 400,000 cells/mL) was seeded per well using a multichannel repeater pipette. When tested, Sypro dyes were dispensed using method 4. PCR plates containing cells, test compound, and BRETSA™ Probe were capped with a plastic lid (Nunc Microplate lid, ThermoFisher Scientific), mixed on an orbital shaker for 15 s at 800rpm, and incubated for 1 hr (unless otherwise noted) in a humidified 37°C/5% CO_2_ incubator. Following equilibration, the plates were removed from the incubator, the lid was removed and the plate was sealed with an aluminum adhesive plate seal. Sealed plates were subjected to thermal challenge using a gradient thermal cycler (Bio-Rad C1000 touch, 96-well format). Thermal challenges were executed for 3 min. Unless otherwise noted, thermal challenges were either gradients from 39°C–60°C (live-cell range), or a single temperature (isothermal, chosen as a balance between isothermal engagement window and sensitivity to potency and often near the EC_80_ of the melt curve). After thermal challenge, the thermocycler protocol was used to rapidly return the plate to 21°C after which TarSeer™ Detection Substrate (Promega) and TarSeer™ Debris Quencher (Promega) were added according to the manufacturer’s recommended protocol. Filtered luminescence was measured on a GloMax^®^ Discover luminometer equipped with 450nm BP filter (donor) and 600nm LP filter (acceptor), using 0.5 s integration time. Data processing including calculation of BRET ratios, BRET value normalization, and curve fitting for melt temperature and compound potency was identical to that described above for the 384-well format.

#### BRETSAs for p53(Y220C)

For understable protein variants such as p53(Y220C), the transfection and assay protocol was carried out similarly to that described above for the 384-well BRETSA™ protocol, with the following modifications. After combining the transfection complex with HEK-293 cell suspension in a cell-culture flask, the cells were incubated for 6hr in a 37°C 5% CO_2_ incubator, after which the cells were transferred to 24°C 5% CO_2_ incubator for the remainder of the transfection. After harvesting the cells and dispensing test compound and BRETSA™ Probe 01, the plates were equilibrated for 60 min in a 24°C 5% CO_2_ incubator. Finally, the thermal gradient was from 30– 54°C. All other aspects of the protocol were unchanged.

#### BRETSAs at elevated temperatures (> 60°C)

When BRETSA™ protocols were executed for highly thermostable proteins (RAS, GTPases and WDR5), BRETSA™ was carried out either using a thermal gradient from 56°C–80°C or at a single isothermal temperature, the StabiLuc™ luciferase was used as the fusion reporter, and the debris quencher was omitted from the detection solution. All other aspects were similar to that described above for 384-well or 96-well formats.

#### BRETSAs in Permeabilized Mode

Permeabilized format BRETSA™ protocols were executed as described above, except that cells were treated with digitonin at a final concentration of 50µg/mL, equilibrations with BRETSA™ Probe 01 and test compound were for 30 min at RT, and the debris quencher was omitted from the detection reagent.

#### Isothermal engagement window analysis

The fold change in the BRET signal induced by a test compound (termed the isothermal engagement window) was used as both a comparative metric between candidate denaturation-sensitive probes (section 1, *vide supra*) as well as a simple and empirically determined criterion for hit-calling (section 2, *vide supra*). For stabilizers or antagonists of the BRET signal, isothermal engagement window is calculated at a specific temperature by dividing the BRET value for the vehicle-treated sample by the BRET value for the compound treated sample to produce a ratio > 1. For destabilizers or agonists of the BRET signal, isothermal engagement window is calculated at a specific temperature by dividing the BRET value for the compound-treated sample by the BRET value for the vehicle-treated sample to produce a ratio > 1. A hit cutoff of isothermal engagement window ≥1.3-fold was established empirically using 146 known target–compound interactions **[Extended Data Table 1].** Ninety-seven percent (142/146) of these previously validated interactions produced isothermal engagement windows ≥1.3-fold **[Extended Data Table 1 and Supplementary Data Set 1]** with isothermal dose response behavior **[Supplementary Data Set 2]** that fit well to sigmoidal binding models (e.g. Equation 3) above. This simple and empirical hit-calling metric agreed well with a statistically defined area-under-the-curve analysis (*vide infra*).

#### Area-under-the-curve (AUC) analysis

Melt-curve data were analyzed with a custom Python script that quantified each compound’s effect on target thermal stability as the area between its melt curve and that of the matched vehicle control. Each compound melt curve was paired with the corresponding DMSO melt curve for the same target within the same biological replicate, and the analysis was restricted to temperatures at which both curves carried a measurement. At each shared temperature the difference between the curves was taken as:

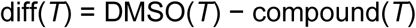

Where a positive difference corresponds to stabilization of the target by the compound and a negative difference to destabilization. The difference curve was divided into contiguous regions of constant sign, and each region was integrated by the trapezoidal rule. Region boundaries were extended on either side to the temperature at which the difference crossed zero, obtained by linear interpolation between the flanking measurements, so that area spanning a change in sign was assigned to the adjacent regions rather than being lost. Stabilizing and destabilizing areas were summed separately as absolute magnitudes. To limit the contribution of spurious noise, only regions supported by at least 2 consecutive measured temperatures of the same sign (stabilizing or destabilizing) were included in AUC calculations; interpolated crossings did not count toward this length and rejected regions contributed no area. Finally, each area was normalized to the integral of the corresponding DMSO curve over the same temperatures and expressed as a percentage, so that compound effects are reported relative to the total signal available to change.

To establish a hit-calling threshold for %AUC, the average technical noise was quantified from 218 independent 384-well melt curve datasets (142 target constructs, with one or two biological replicates each). For each melt curve, the DMSO wells of the two technical replicates were treated as a compound-vehicle pair and the total absolute %AUC (irrespective of direction) was calculated without the consecutive run-length filter described above. This measure captures all variation between curves that should be identical, and it is an upper bound on the filtered, single-direction values used for hit calling. The resulting values were positively skewed and were cube-root transformed to approximate a normal distribution (transformed mean, 1.54; standard deviation, 0.18). A threshold of 10 %AUC corresponded to approximately 3.4 standard deviations above the mean noise level (95% bootstrap interval, 3.1 to 3.8), and none of the 218 noise values reached this level (maximum, 9.6 %AUC). A threshold of 10 %AUC was therefore adopted for hit calling, and its application is described for each assay below. The same threshold was applied to 96-well data, for which too few DMSO comparisons were available to estimate noise separately.

#### AUC analysis for Broad Target Survey (Related to *Section 2*)

Melt curves for 114 full-length target-ligand pairs were analyzed with the AUC method described above. Technical replicates were averaged, and each biological replicate was analyzed separately. A replicate was scored as a hit if the stabilizing or destabilizing %AUC was at least 10%. A target-ligand pair was called a hit when this criterion was met in both biological replicates, or in the single replicate when only one was collected.

#### Longitudinal BRETSA™ Assays and Area Under Curve (AUC) Measurements

Longitudinal BRETSA™ assays were performed in 384-well format using in-well transfection, in which DNA, transfection reagent, and cells were combined directly in the PCR assay plate in place of the flask-based transfection described above. HEK-293 cells were dissociated as described above. Cells were pelleted at 200 × g for 3 min and resuspended in Opti-MEM (Gibco) containing 2% FBS to a final density of 1 × 10^6^ cells/mL. Plasmids encoding NanoLuc^®^ fusion proteins were diluted into Transfection Carrier DNA (Promega) at a 1:9 (mass/mass) ratio and concentration of 20µg/mL. For 300 kinase targets, the optimal NanoLuc^®^ fusion orientation was selected using prior knowledge from existing NanoBRET^®^ Target Engagement assays (Promega). For the remaining targets without a validated NanoBRET^®^ TE assay, both fusion orientations were tested where available. Using an Agilent Bravo liquid handler, 5µL of pre-mixed DNA was stamped into assay plates. NanoLuc^®^ control wells (unfused NanoLuc^®^ with Transfection Carrier DNA) and transfection control wells (no DNA, TE buffer only) were included in each experiment. FuGENE HD (Promega) was diluted to 0.06U/µL in Opti-MEM, and 5µL was added to each well using a WellJet (Integra). Complexes were allowed to form for 15 min at room temperature, after which 10µL of cell suspension was dispensed to each well using the WellJet (10,000 cells/well; final FBS 1%). Plates were briefly centrifuged to collect the contents and incubated for 20−24 hr in a humidified 37°C/5% CO_2_ incubator.

Following incubation, BRETSA™ Probe-01 (Promega) was dispensed directly from DMSO to a final concentration of 1x using a droplet digital dispenser (Tecan D300e). Plates were shaken for 5 s on the dispenser and then mixed on a BioShake (Q Instruments) at 3000rpm for 30 s. Staurosporine (Selleckchem) was then dispensed directly from DMSO to a final concentration of 10µM, alongside vehicle control wells, and DMSO was normalized across all wells. Another 5 s shake was performed on the dispenser. Plates were incubated for 1 hr in a humidified 37°C/5% CO_2_ incubator prior to heat treatment. Then, 10µL of 3x TarSeer™ Detection Substrate (Promega) and TarSeer™ Debris Quencher (Promega) was added to each well using the WellJet. Plates were sealed with an optically clear adhesive microplate seal (Bio-Rad Microseal ‘B’) and subjected to iterative thermal challenge using a Bio-Rad PTC Tempo thermal cycler, with the lid temperature set 5°C above the block temperature. Plates were held for 30 s at each temperature from 37°C to 59°C in 2°C increments. After each temperature challenge, plates were cooled to 21°C, loaded into a 384-well cross-talk reduction plate adapter (Promega), and read on a PHERAstar FSX (BMG Labtech). After reading, the plate was returned to the thermal cycler for the next heat challenge step. Filtered luminescence was measured using a 450/80nm BP filter (donor) and a 610nm LP filter (acceptor), with a 0.2 s integration time, a 14.5mm focal height, and a gain of 2000 for both channels. BRET ratios were calculated as described above. Prior to melt-curve analysis, targets were filtered on NanoLuc^®^ donor signal to remove wells with failed transfections or poorly expressing targets. Targets were excluded if their DMSO donor signal at 37°C did not exceed 7σ above the untransfected control well signal. Each target was measured in technical singlicate per condition, and biological duplicate.

Melt midpoints (apparent T_M_) were obtained by fitting a 4PL curve to the vehicle-treated curve for each target in JMP software (version 18). Because the applied gradient spanned 37–59°C, a T_M_ was assigned only where the two biological replicates were concordant: where both returned a numeric midpoint within 10°C of one another, the mean is reported; where both exceeded 60 °C, the target is reported as >60°C, these values lying at or beyond the top of the measured gradient. Targets for which one replicate exceeded 60°C while the other returned a numeric midpoint, or for which the two replicates differed by 10°C or more, were not assigned a T_M_. Melt curve data was run through the AUC analysis detailed above, and a 10% threshold was used for hit calling. Only stabilizing compound effect areas were considered. Engagement was assessed independently of melt-curve fitting, since %AUC stabilization integrates the difference between vehicle and compound-treated curves and does not require a fitted midpoint; a small number of engaged targets are therefore reported without a corresponding T_M_. For targets assayed in both NanoLuc® fusion orientations, a single construct was selected for reporting: where both orientations yielded analyzable data, the construct with NanoLuc® nearer the annotated kinase domain was preferred, with melt-curve quality and replicate agreement used to resolve equivocal cases. In a small number of cases the orientation giving the better-resolved melt differed from that giving the reproducible engagement call, and the two measurements are reported from their respective constructs.

#### BRETSA Assays for Ternary Complex Cooperativity

To determine cellular ternary complex cooperativity for HLDA-221 or KRAS Tricomplex molecular glues, transfections were performed as described above except that DNA ratios in the transfection complex were 1 part StabiLuc™ (or NanoLuc^®^) fusion vector to 4.5 parts untagged partner protein expression vector (i.e. FKBP12(F36V) or CypA), and 4.5 parts transfection carrier DNA. BRETSA™ assays were performed as described above in the 96-well format with dispensing of BRETSA™ Probe 01 at the 1X dose using method 3. Experiments with HLDA-221 and related compounds were performed in HEK-293 cells. Experiments with the KRAS Tricomplex molecular glues were performed in HAP1 CypA(–) cells.

#### NanoBRET^®^ Target Engagement Assays

NanoBRET^®^ assays for HDAC1, CRBN/DDB1, and PARP1 were executed as described in the respective manufacturer’s recommended protocols (Promega). Briefly, for HDAC1-NanoLuc and NanoLuc-CRBN/DDB1, cells were transfected as described above in bulk in T225 flasks, after which they were harvested by trypsinization as described for the BRETSA™ assays, resuspended in Opti-MEM medium (without phenol red), and seeded into 96-well Corning^®^ non-binding surface plates (Cat.# 3600) at 20,000 cells per well. NanoBRET^®^ Tracers (Promega) for HDAC1 and CRBN were prepared at a working concentration of 20X in tracer dilution buffer (Promega) at a concentration of 20µM or 10µM, respectively. Test compounds were prepared at a working concentration of 10X in Opti-MEM (without phenol red). Plates containing transfected cell suspension (85µL) were treated with 5µL 20X NanoBRET^®^ Tracer and 10µL 10X test compound, after which they were mixed on an orbital shaker for 15 s at 800rpm.

For PARP1-NanoLuc, cells (5,000 per well) were transfected as described above directly into 384-well Corning^®^ tissue culture-treated assay plates (Cat.# 3570) in Opti-MEM medium (without phenol red) containing 1% FBS. NanoBRET^®^ Tracer PARP-01 (Promega) was dispensed directly from DMSO to a final concentration of 12.5nM in the assay. Test compounds were dispensed directly from concentrated DMSO stocks into the assay plates using an acoustic dispenser (Beckman Echo 650).

Plates containing transfected cells, NanoBRET^®^ tracers, and test compounds were incubated for 2 hr in a humidified 37°C/5% CO_2_ incubator. Following equilibration, plates were removed from the incubator and cooled to room temperature, after which NanoBRET^®^ Target Engagement Substrate (Promega) and Extracellular NanoLuc^®^ Inhibitor (Promega) were added according to the manufacturer’s recommended protocol. Filtered luminescence was measured as described above for BRETSA™ assays. Data processing including calculation of BRET ratios and BRET value normalization, and curve fitting for compound potency was identical to that described above for BRETSA™ assays.

Cellular kinase selectivity for staurosporine (10µM dose) was profiled using the NanoBRET^®^ TE Kinase Selectivity platform (K300 panel, Promega) against a panel of 300 full length kinases^25-27^.

#### Cell Viability Measurements in the presence of BRETSA™ Probe 01 or Sypro Dyes

The impact of BRETSA™ Probe 01 or Sypro dye treatment on cell viability was determined using the CellTiter-Glo^®^ 2.0 assay (Promega) according to the manufacturer’s recommended protocol. Briefly, impact to viability was assessed in both a dose- and time-dependent manner. HEK-293 cells were harvested by trypsinization and seeded into Corning^®^ 96-well plates (Cat.# 3600) at 40,000 cells per well in Opti-MEM medium without phenol red, 100µL volume cell suspension per well. BRETSA™ Probe 01, Sypro Orange, or Sypro Red were dispensed to the plates directly from their concentrated DMSO stock solutions using a Tecan D300e to final concentrations as noted in the corresponding figures and legends. DMSO was included as the vehicle-treated control and digitonin was included at a final concentration of 50µg/mL as a cell death inducing control. After probe dispense, plates were immediately mixed at 700rpm for 15 s to ensure probe dispersal, after which they were incubated for fixed periods of time (as indicated) in a 37°C/5% CO_2_ incubator. Subsequently, plates were cooled back to ambient temperature and CellTiter-Glo^®^ 2.0 reagent was added, after which the plates were mixed on an orbital shaker for 2 min at 500rpm and then incubated for another 8 min at ambient temperature. Total luminescence was measured on a Glomax^®^ Discover multi-mode reader using the CellTiter-Glo^®^ protocol with the default settings. Apparent viability was reported as the relative luminescence (%) compared to time zero controls collected for each sample type.

#### Differential Scanning Fluorimetry (DSF)

Differential scanning fluorimetry experiments were carried out for PARP1, MAPK14, AURKB, ABL1, and HSP90AB1 using Sypro Orange dye (Thermo Fisher). Briefly, a master mix of the protein of study was prepared at a 1.25X concentration in assay buffer (10mM HEPES, 500mM NaCl, pH 7.5) in a 4µL volume, after which it was combined with 1µL of an intermediate solution containing Sypro Orange (35X) and test compound (5X) or DMSO vehicle to produce a reaction (5µL) containing 1X protein, 7X Sypro Orange, and the final concentration of test compound (as indicated). For PARP1, a 1.1µg/µL stock solution of PARP1 protein (Sino biological Cat.# 11040-H08B, full length human PARP1 (NP_001609.2) with a C-terminal polyhistidine tag; provided in 20mM Tris, 300mM NaCl, 10% glycerol, 0.5mM TCEP, 2mM EDTA, pH 7.5) was diluted 4.4-fold into assay buffer to prepare the protein master mix. For MAPK14, protein was tested from 2 different sources. For MAPK14 source 1, a 0.87µg/µL stock solution of MAPK14 protein (a kind gift from the Stefan Knapp Lab; full length human MAPK14 Uniprot isoform 1 (NP_001306.1) with a T263A mutation; provided in 25mM HEPES, 500mM NaCl, pH 7.5) was diluted 4.6-fold into assay buffer to prepare the protein master mix. For MAPK14 source 2, a 0.3µg/µL stock solution of MAPK14 protein (Sino biological, Cat.# 10081-H07B; full length human MAPK14 (NP_001306.1) with a N-terminal polyhistidine tag; provided in 20mM Tris, 500mM NaCl, 10% glycerol, pH 7.4) was diluted 1.6-fold into assay buffer to prepare the protein master mix. For AURKB, a 1.4µg/µL stock solution of AURKB protein (a kind gift from the Stefan Knapp Lab; Uniprot isoform 1 of human AURKB(55-345, M298T); prepared in 25 mM HEPES, 500 mM NaCl, pH 7.5) was diluted 3.2-fold into assay buffer to prepare the protein master mix. For ABL1, a 0.66µg/µL stock solution of ABL1 protein (a kind gift from the Stefan Knapp Lab, Uniprot isoform 1 of human ABL1(229-512); prepared in 25mM HEPES, 500mM NaCl, pH 7.5) was diluted 3.2-fold into assay buffer to prepare the protein master mix. In the final reaction mixtures, PARP1, MAPK14 (source 1), MAPK14 (source 2), AURKB, and ABL1, proteins were present at final concentrations of 1.7, 3.5, 3.7, 10.4, and 5.0µM, respectively. Reactions were assembled as described above in 96-well white PCR plates (Bio-Rad HSP9655) sealed with Bio-Rad Microseal ‘B’ optical seal (Cat.# MSB1001) and were pre-equilibrated at 25°C for 10 min. Following equilibration, thermal cycling and fluorescence measurements were carried out using a Bio-Rad CFX96 thermocycler with the following settings: thermal gradient 25–95°C, increment 0.5°C, ramp rate 2°C/s, equilibration time 15s, lid temperature 95°C, excitation filter 515–535nm, and emission filter 560–580nm. RFU data at each temperature, as well as first-derivative (-dF/dt) data were exported from the CFX Maestro software (version 5.3.022.1030) and re-plotted in GraphPad Prism 11. Melt temperatures (T_M_) were determined using the first-derivative minimum of –dF/dT. DSF dose response curves were constructed by plotting the apparent T_M_ values as a function of compound dose in GraphPad Prism 11.

#### BiTSA isothermal screening assays

The HiBiT tag was endogenously engineered onto the N-terminus of BRD4 in HEK-293 cells by CRISPR/Cas9. BRD4 BiTSA deployed at screening scale followed a published protocol^32^. Briefly, dose responses of test compounds were dispensed into empty 384w PCR plates using an Echo655 acoustic dispenser, and 10,000 HiBiT-tagged BRD4 HEK-293 cells were seeded per well as 10µL in DMEM media (Gibco) supplemented with 1% FBS, using a Multidrop Combi dispenser, then sealed using a PlateLoc. Following 1 hr incubation at RT, a 2 min heat shock was applied at indicated temperature using an On Deck Thermal Cycler (ODTC, Inheco), and plates were briefly spun then de-sealed using an XPeel before 10µL Nano-Glo® HiBiT Lytic Detection reagent (Promega N3030) diluted 1:1 in PBS was added using a Certus Flex (0.2/0.1mm valve). Lysates were mixed by 1 min at 2500rpm on a BioShake 1000, incubated 5 min RT, then centrifuged at 400 x g 1 min. 10µL of resulting lysates was transferred to a Greiner 784075 detection plate using a PRIME liquid handler (HighRes), and luminescence was analyzed from the detection plate using a PheraStar FSX plate reader.

Cell models with the HiBiT tag endogenously engineered onto the N-terminus of DNPH1 in SUM149PT cells by CRISPR/Cas9, DNPH1 BiTSA experiments, and DNPH1 FRET binding assay experiments, were performed as described as previously reported^33^. Briefly for DNPH1 BiTSA, dose responses of test compounds were dispensed into empty 384w PCR plates using an Echo655 acoustic dispenser, and 5,000 HiBiT-tagged DNPH1 SUM149 cells were seeded per well as 10µL in Nutrient F12 Ham (Sigma N6658) supplemented with 1% FBS, using a Certus Flex, then manually sealed. Following 1h incubation at RT, a 2 min heat shock was applied at indicated temperature using an ODTC, and plates were manually de-sealed, before 10µL Nano-Glo® HiBiT Lytic Detection reagent diluted 1:1 in PBS was added by Certus Flex. Lysates were mixed by 1 min Multidrop shake, incubated 5 min at RT, then centrifuged at 300 x g 1 min. 10µL of resulting lysates were transferred to a Greiner 781080 detection plate using a Bravo liquid handler (Agilent), and luminescence was analyzed from the detection plate using a PheraStar FSX plate reader.

Assay robustness during assay development and on screening plates was calculated as the Robust Z’ Factor (RZ’) between Max effect (30µM control Inhibitor) and Min effect (DMSO control) treated samples via the equation:

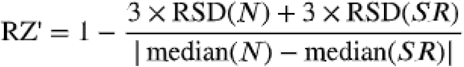

Where RSD is the Robust Standard Deviation, N is the neutral DMSO control wells, and SR is the Inhibitor control wells.

#### BRETSA™ isothermal screening assays for comparison to BiTSA (BRD4 and DNPH1)

BRETSA™ screening assays deployed at screening scale followed a similar protocol as reported above. StabiLuc-BRD4 and DNPH1-StabiLuc™ constructs were generated in pSB constructs under a CMV promoter with flexible VSLGSSG linkers between the ORF and the StabiLuc™ ORF (vector details available on request). HEK-293T cells were transfected 18 hr with scaled transfection ratios; 3µg DNA, 27µg Transfection carrier DNA, 90µL FuGENE HD in 3mL OptiMEM (Gibco), mixed with 20 parts (60 mL) HEK-293T cells suspended at a density of 2 x 10^5^ per mL in DMEM + 10% FBS. Harvested cells were resuspended to 4.5 x 10^5^ cells/mL in DMEM + 1% FBS and BRETSA™ Probe 01 was added directly to the cell suspension (method 2). BRETSA™ Probe 01 was added 1:1000 to StabiLuc-BRD4 cells (1X) and 1:500 to DNPH1-StabiLuc™ cells (2X). 20µL of cell/probe suspension was seeded per well into assay ready 384w PCR plates (BioRad) containing test compounds as above using a Multidrop Combi with a small metal tip cassette primed via seeding a dummy plate first (BRD4) or using a Dragonfly dispenser (DNPH1). For larger screening runs, cell/probe suspension were stirred via magnetic stir bar at 250rpm in a polystyrene container between seeding plates. Plates were sealed and incubated 1h RT, followed by a 2 min heat shock applied at indicated temperature using a PTC Tempo PCR machine (BRD4) or an ODTC heating block (DNPH1). Plates were cooled at RT for 2–3 min. TarSeer™ Detection Substrate (Promega) and TarSeer™ Debris Quencher (Promega) were prepared in HBSS according to manufacturer’s recommended protocol and added 10µL/well by either Certus Flex (BRD4) or Dragonfly dispenser (DNPH1). For DNPH1 experiments, the Debris Quencher was omitted. Following a 10 min incubation, assay plates were placed into the 384-well plate adaptor (Promega Cat #GM3055) and NanoBRET^®^ emissions measured on a PheraStar FSX with optic modules LUM 610-LP 450-80 and focal height set to 11.5mm. NanoBRET^®^ mBU was calculated as above, or in screening plates calculated as % effect, the ratio of acceptor (610nm) / donor (450nm) for the entire plate was normalized to on-plate Inhibitor control and DMSO control (n = 12 each) signals in Genedata Screener via the equation:

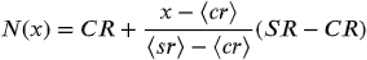

Where x is the measured raw signal value of a well, N(x) is the normalized value of the raw signal, <CR> is the median of the measured signal values for the DMSO controls, <SR> is the median of the measured signal values for the Inhibitor controls, CR is the desired median normalized value for the DMSO controls, and SR is the desired median normalized value for the Inhibitor controls.

## ACKNOWLEDGEMENTS

The authors thank Kristin Huwiler, Tom Livelli, Andrew Zhang, Frank Fan, Kelly Teske, and Jennifer Wilkinson for thoughtful discussions about the research and data interpretation, and Hyojin Kim and Sabrina Winfield for aiding construct design. Structural Genomics Consortium is a registered charity (no: 1097737) that receives funds from Bayer AG, Boehringer Ingelheim, Bristol Myers Squibb, Genentech, Genome Canada through Ontario Genomics Institute [OGI-196], Canada Foundation for Innovation Ontario Research Fund, MITACS, EU/EFPIA/OICR/McGill/KTH/Diamond Innovative Medicines Initiative 2 Joint Undertaking [EUbOPEN grant 875510], Janssen, Merck KGaA (aka EMD in Canada and US), Pfizer and Takeda.

## AUTHOR INFORMATION

### Author Contributions

J.D.V, J.S., A.M., D.B.-L., W.Z, M.C, P.L.M., and M.B.R. guided the study and supervised research from their respective groups. M.B.R, J.D.V, A.M., J.S., A.S., and M.T.B. wrote the manuscript. S.R., and C.A. designed and prepared expression constructs. I.H, M.Z., and C.R.C. Provided BRETSA™ Probe 01. P.S., J.D.V, M.T.B., A.M., A.S, C.H., M.P.S., B.G., K.N., B.M., K.K.D.-H., T.B.L., R.L.J., M.R.S., C.A., J.S., S.B., L.U., C.F., M.V.-C., M.M.S., and M.B.R. executed BRETSA™ experiments. T.B.L. executed DSF experiments. J.S., S.B., L.U., C.F., and M.V.-C. executed BiTSA experiments. B.G. executed cell viability experiments. J.D.V. executed NanoBRET^®^ TE experiments for correlation to BRETSA™. C.D. executed NanoBRET^®^ TE kinase selectivity profiling experiments for staurosporine

---

**Extended Data Figure 1.**
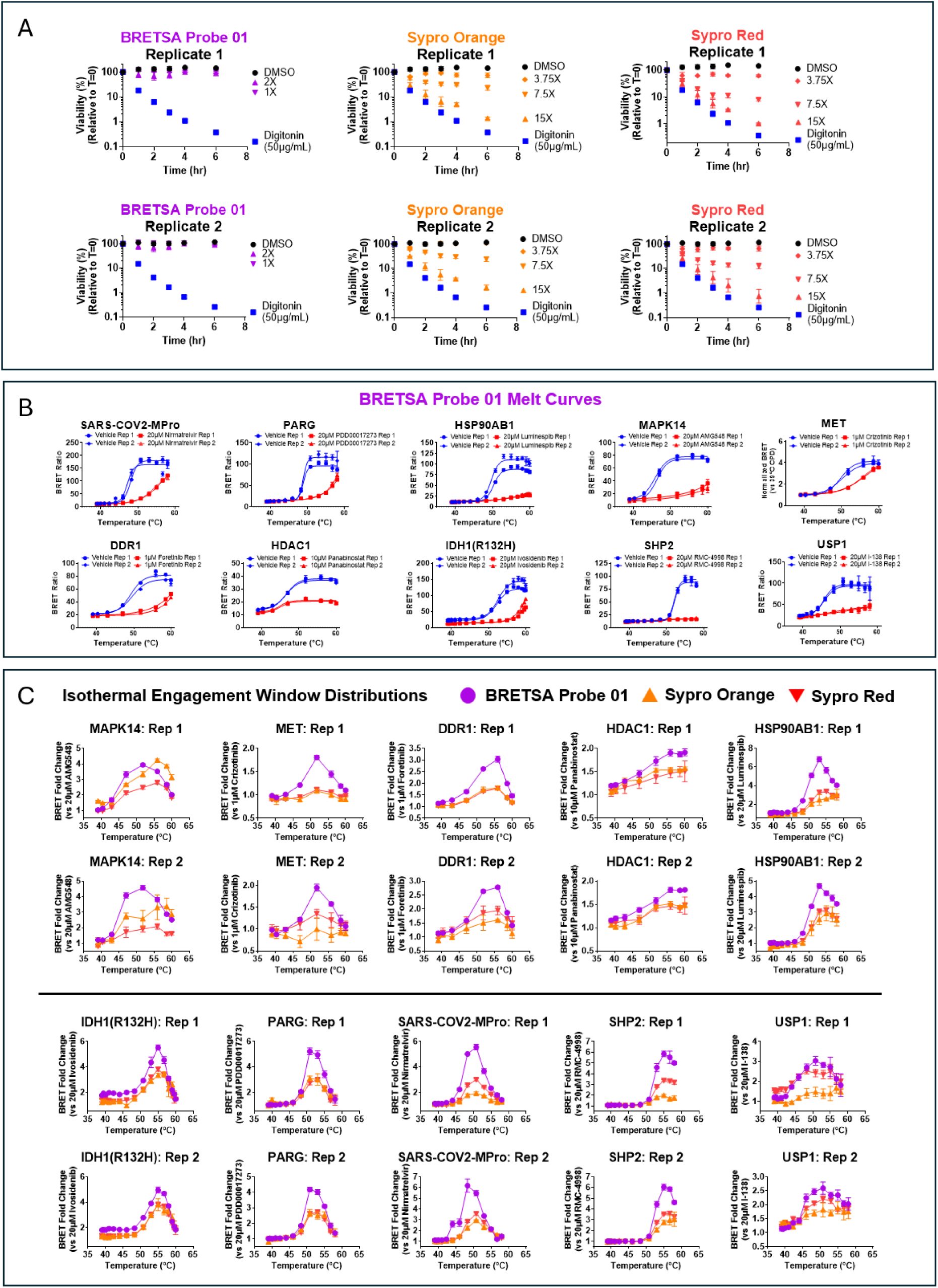
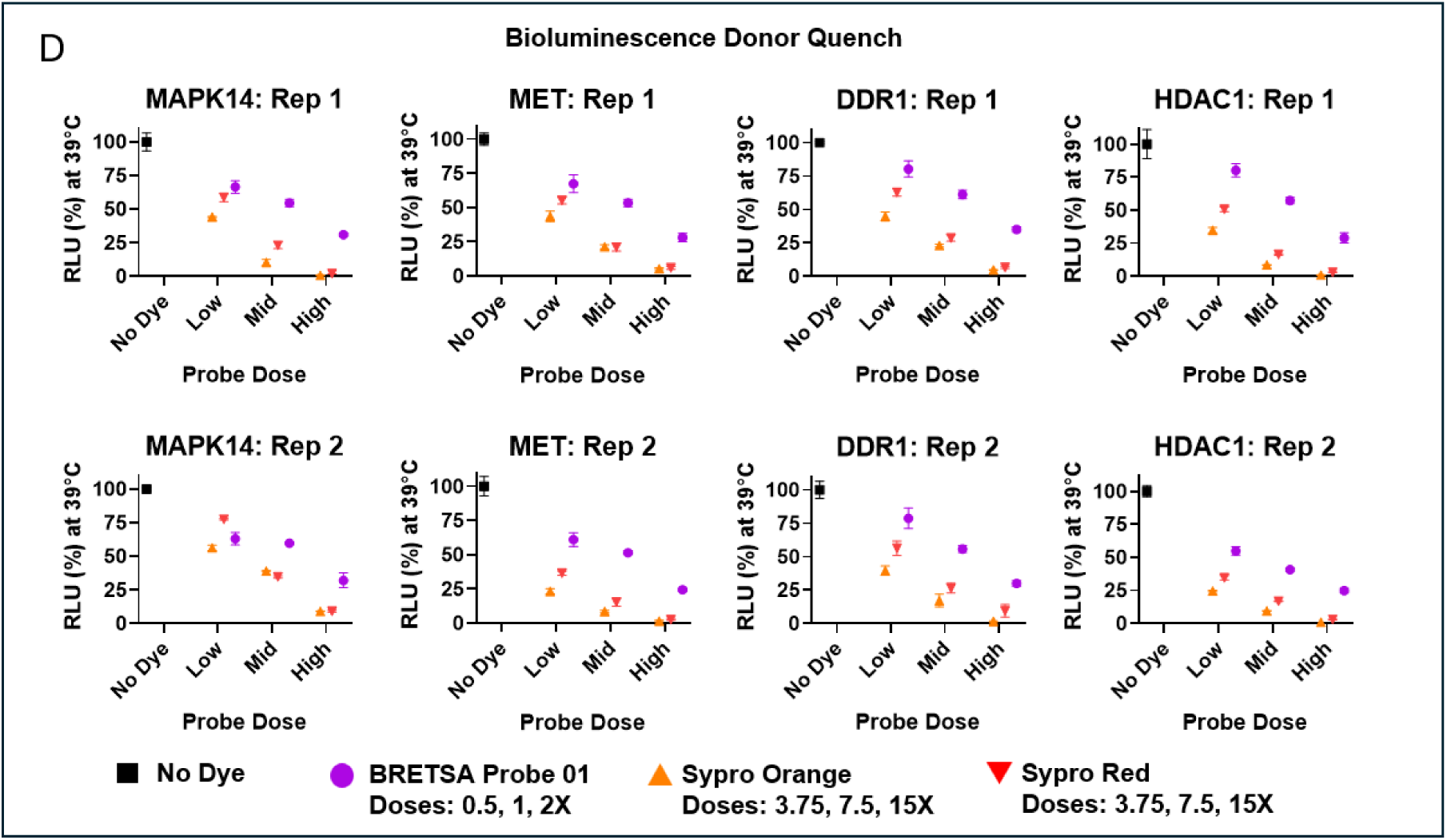
Identifying an ideal probe for the BRET Shift Assay (BRETSA). Related to Fig. 1. (A) HEK-293 cytotoxicity curves biological replicates for BRETSA Probe 01, Sypro Orange, and Sypro Red. Data are from experiments performed twice, where individual data points are the mean (± SD) of 4–8 technical replicates. (B) Biological replicates for BRETSA melt curves using BRETSA Probe 01 for 10 sentinel target protein-luciferase fusions expressed inside live HEK-293 cells and treated with a saturating dose of a known cell permeable ligand. Data are from experiments performed twice, where individual data points are the mean (± SD) of 3–4 technical replicates. All targets used a 1X dose, except for MET kinase, which used a 2X dose. For any target where the biological replicates were performed on two different readers, the data were normalized to the BRET value of the low temperature compound-treated samples for proper comparison. (C) Biological replicates of isothermal engagement window distributions for the 10 sentinel targets for BRETSA Probe 01, Sypro Orange, and Sypro Red. Isothermal engagement window is calculated by dividing the vehicle-treated BRET ratio by the compound-treated BRET ratio at each temperature. Data are from experiments performed twice where individual data points are the mean (± SD) of 3–4 technical replicates. (D) Dose-dependent luminescence donor quench by Sypro dyes compared to BRETSA Probe 01 at a non-denaturing temperature (39°C) for 4 representative sentinel targets. Data are from experiments performed twice where individual data points are the mean (± SD) of 3 technical replicates.

**Extended Data Figure 2.**
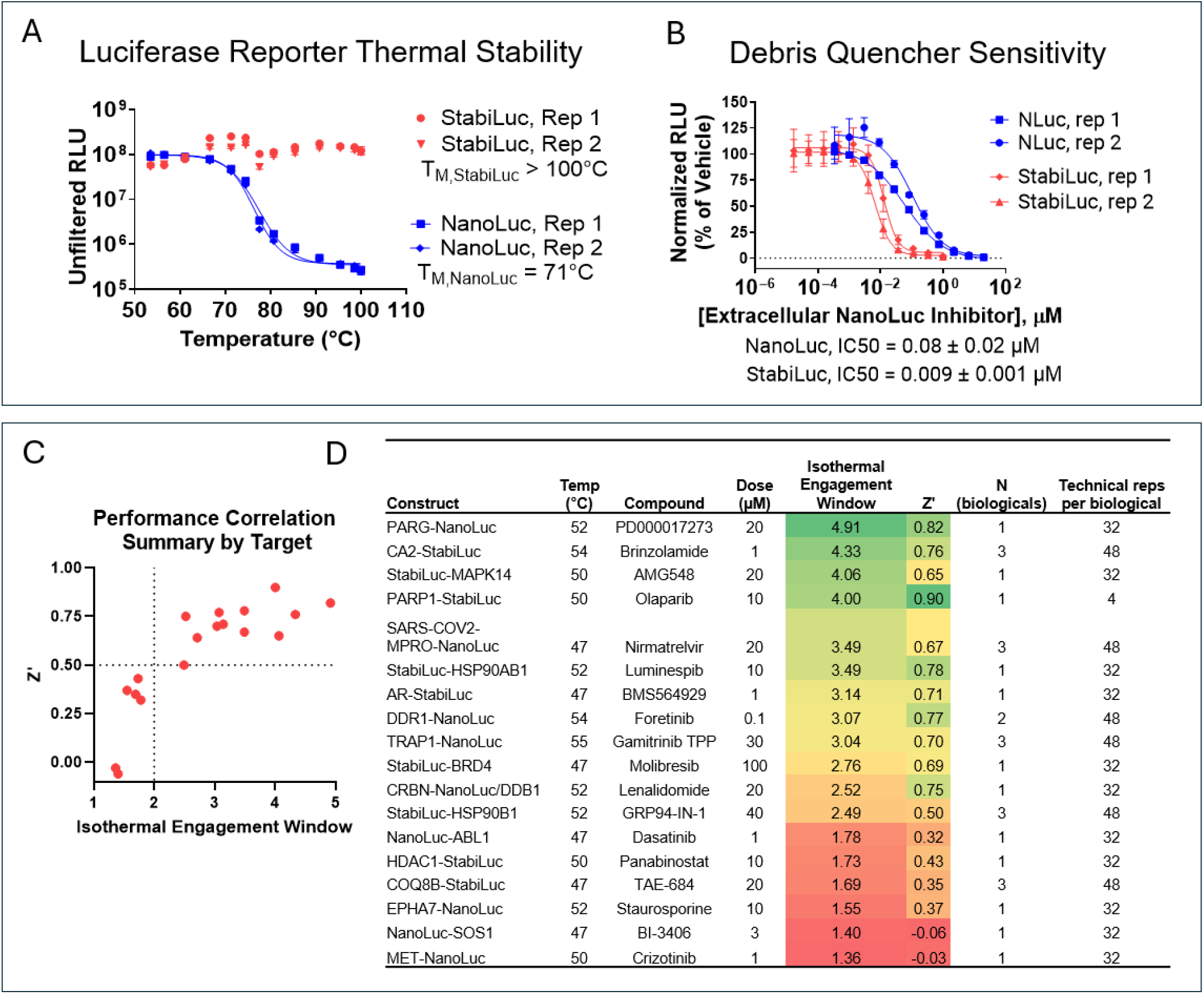
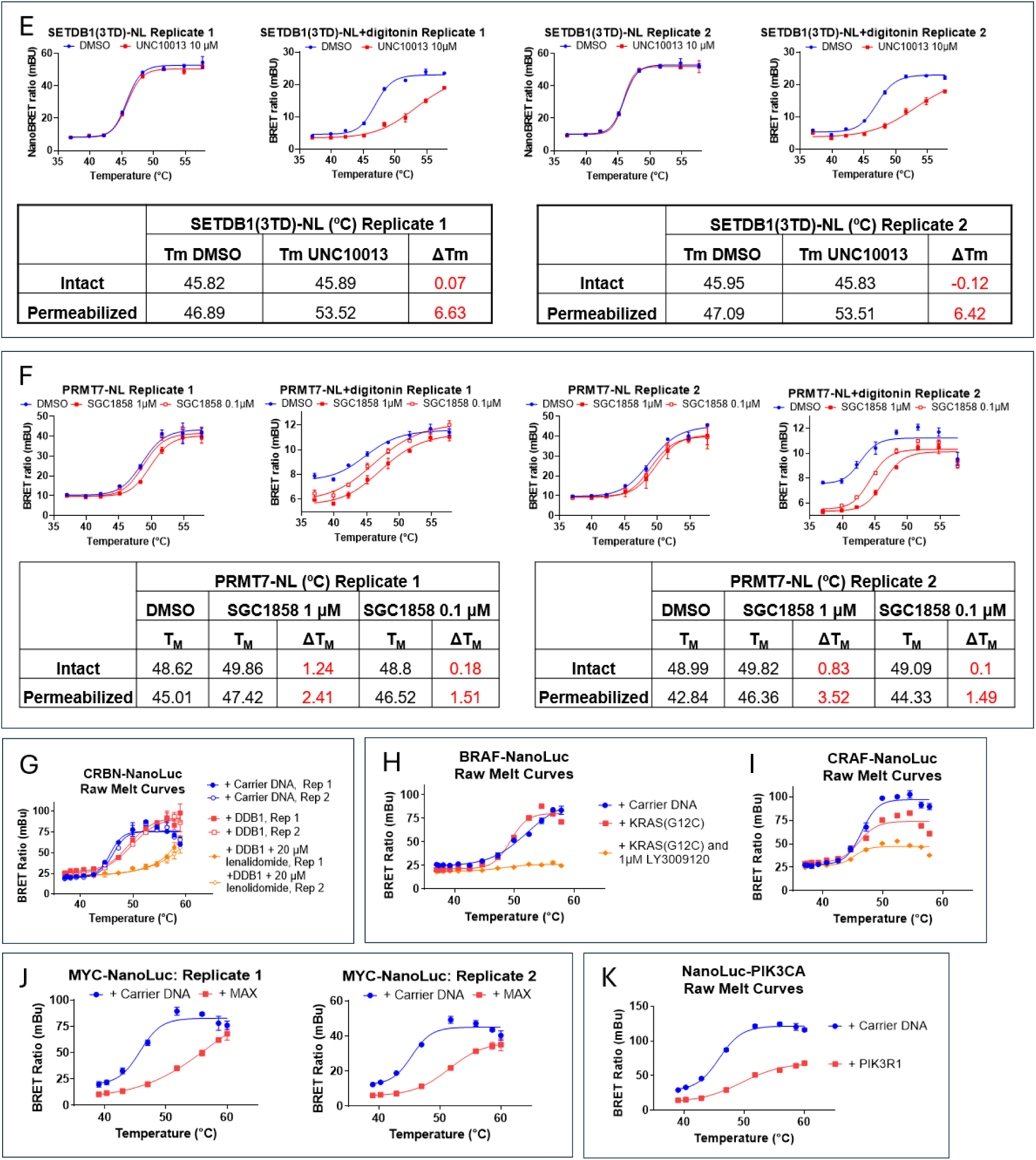
Broad target survey using StabiLuc and NanoLuc reporters. Related to Fig. 2. (A) Biological duplicates for thermal stability of luciferase reporters. Each data point is the mean (± SD) of 5 technical replicates. (B) Biological duplicates of NanoLuc and StabiLuc sensitivity to the Extracellular NanoLuc Inhibitor (Debris Quencher) as measured by titrating the inhibitor into cells expressing either luciferase after digitonin permeabilization. Each data point is the mean (± SD) of 3–4 technical replicates. (C and D) Relationship between isothermal engagement window and assay performance across 18 sentinel BRETSA targets. (C) Individual data points are the mean Z’ and isothermal engagement window for 1–3 independent experiments, each calculated using 4–48 technical replicates. (D) Source data and treatment conditions for the 18 targets plotted in panel C. (E and F) Biological duplicates for comparison of BRETSA melting curves obtained in intact and digitonin-permeabilized cells treated with poorly cell-permeable inhibitors of SETDB1 (E) and PRMT7 (F). Data are from two independent experiments, with individual data points representing the mean (± SD) of 3 technical replicates. (G) Biological duplicates for the influence of DDB1 co-expression and/or lenalidomide treatment on thermal stability of CRBN-NanoLuc using BRETSA. Each data point is the mean (± range) of 2 technical replicates. (H and I) Influence of KRAS(G12C) co-expression and/or treatment with LY-3009120 on the BRETSA melt behavior of BRAF (H) and CRAF (I). Individual data points are the mean of technical duplicates from a single experiment. (J) Biological duplicates of the assessment of the PPI between MAX and MYC-NanoLuc using BRETSA. Each data point is the mean (± SD) of 3–6 technical replicates. (K) Assessment of the PPI between PIK3R1 (p85) and NanoLuc-PIK3CA (p110α) using BRETSA. Each data point is the mean (± SD) of 4 technical replicates from a single experiment.

**Extended Data Figure 3.**
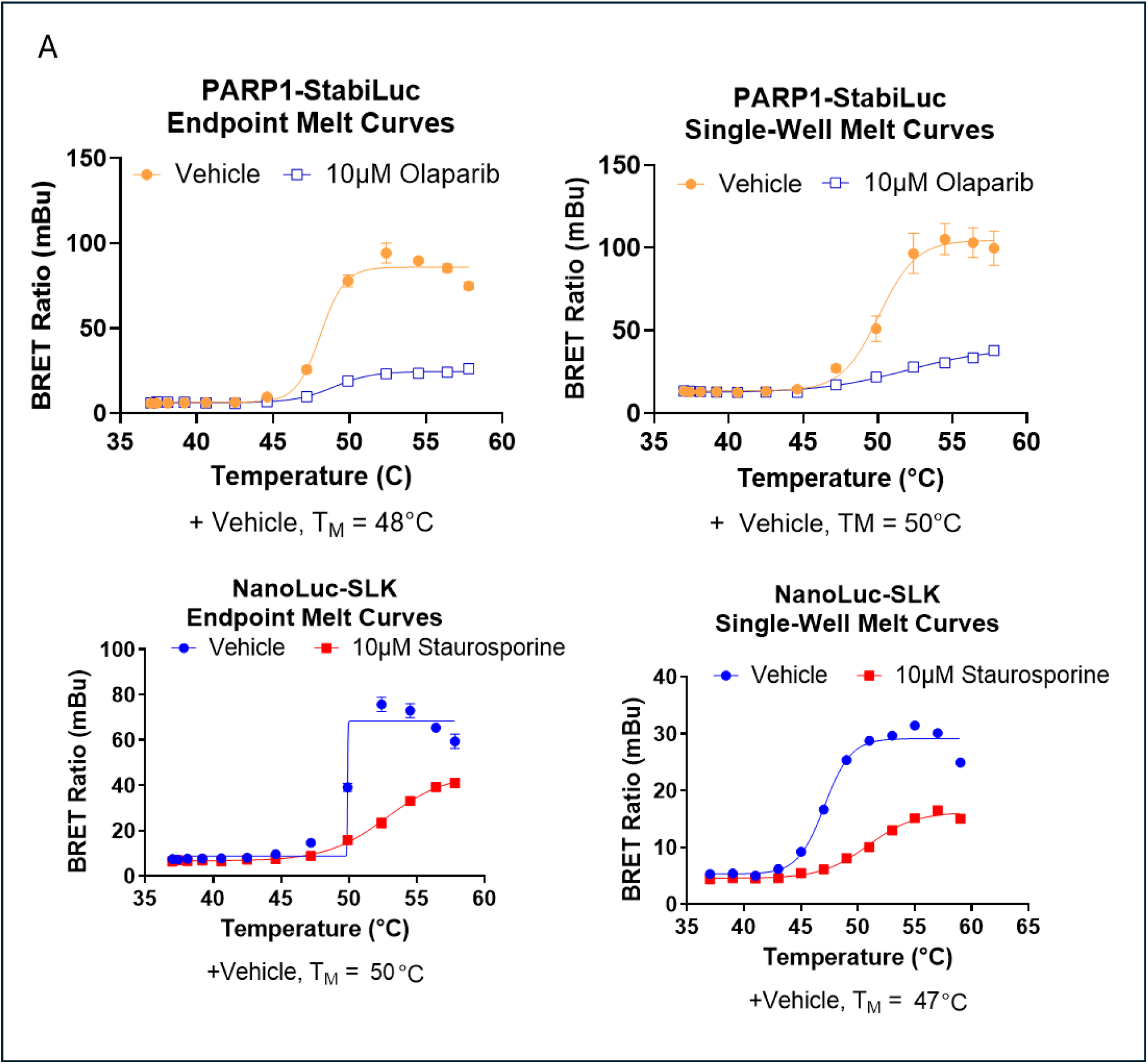
Validation of the longitudinal mode of BRETSA. Related to Fig. 3. (A) Comparison of melt curves obtained via standard endpoint BRETSA and single-well BRETSA for PARP1 and SLK kinase. Experiments were executed as described in the methods. Endpoint data are from a single experiment where individual data points are the mean (± range) of technical duplicates. Single-well data are from a single experiment where individual data points are the mean (± SD) of technical quadruplicates.

**Extended Data Figure 4.**
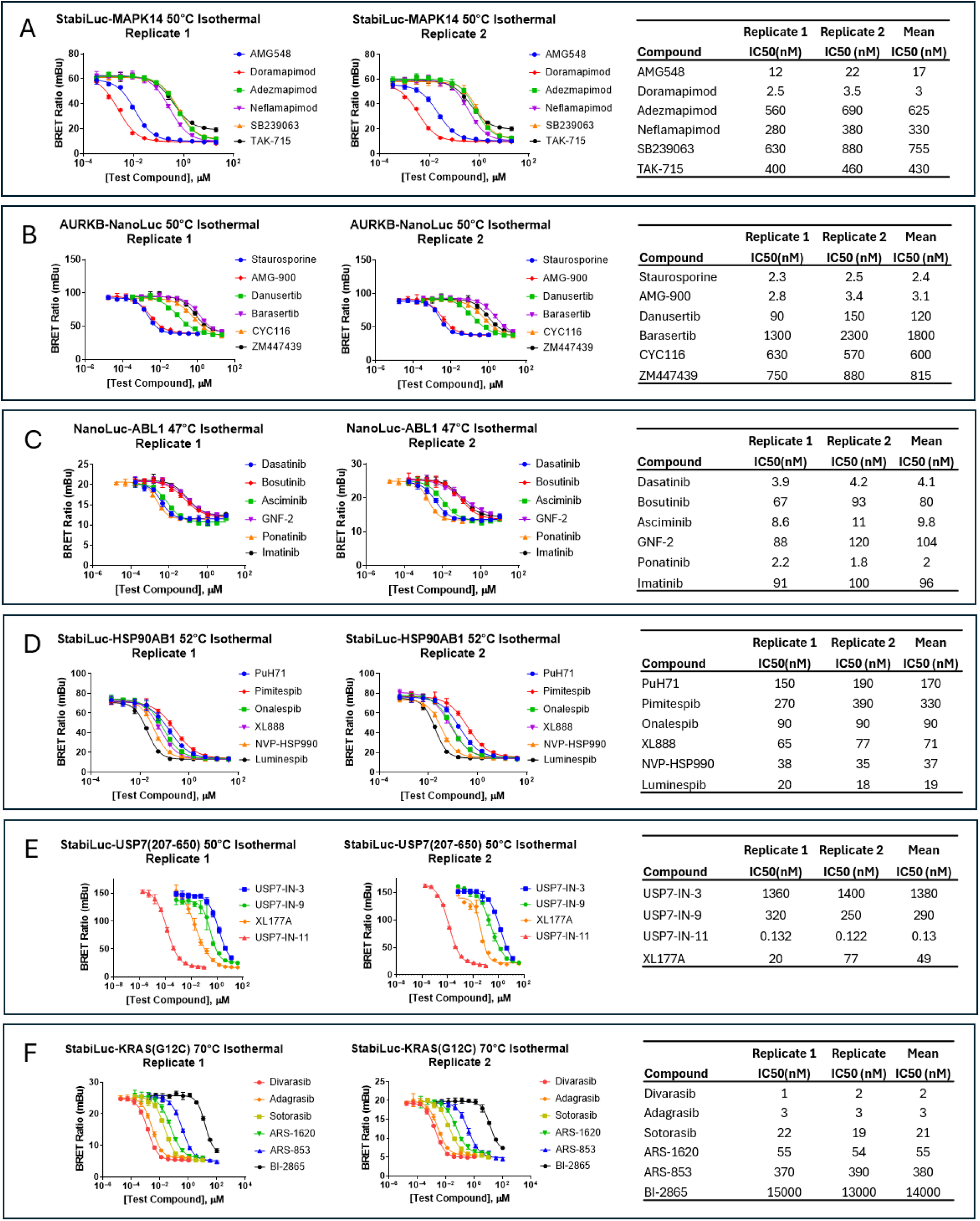
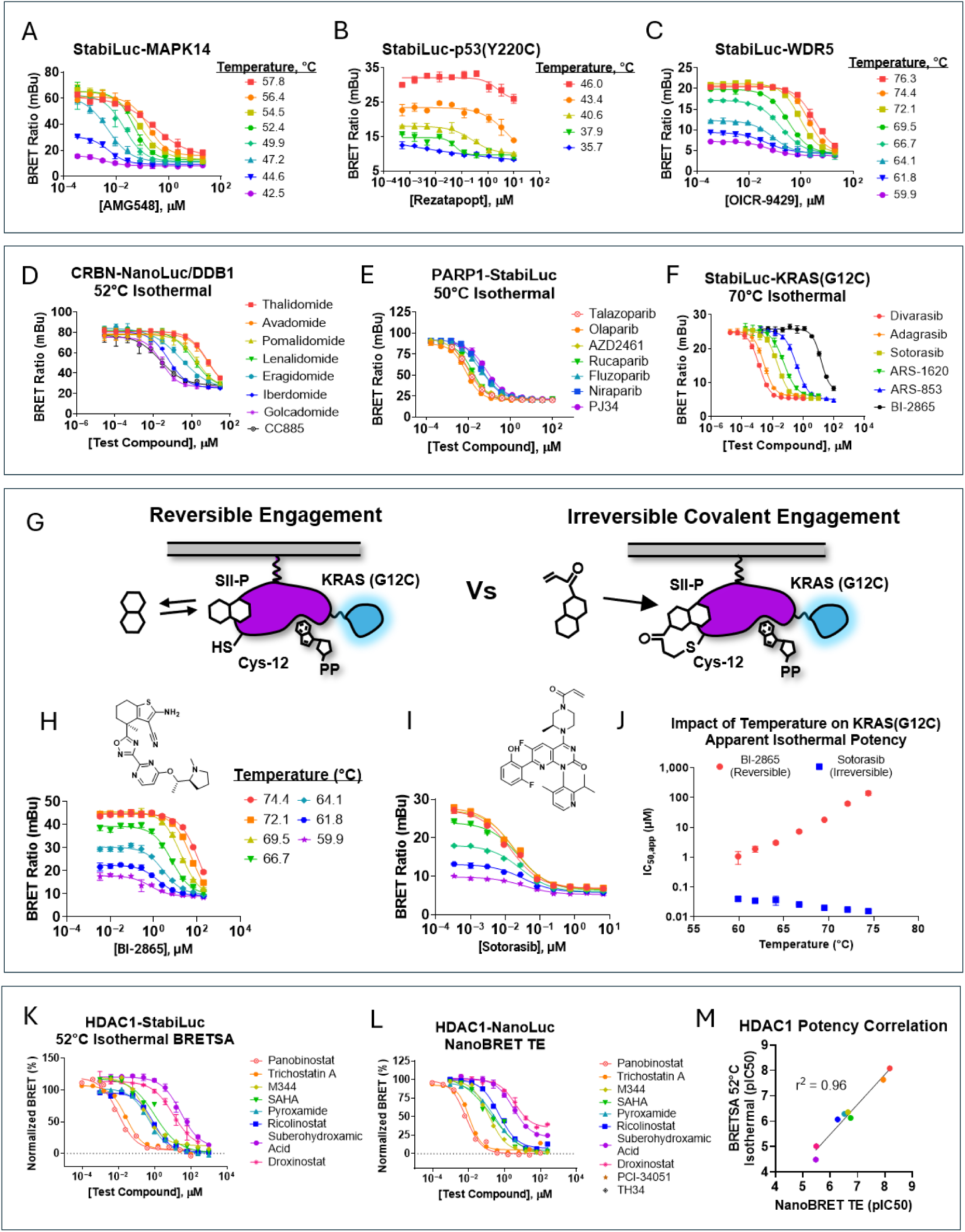
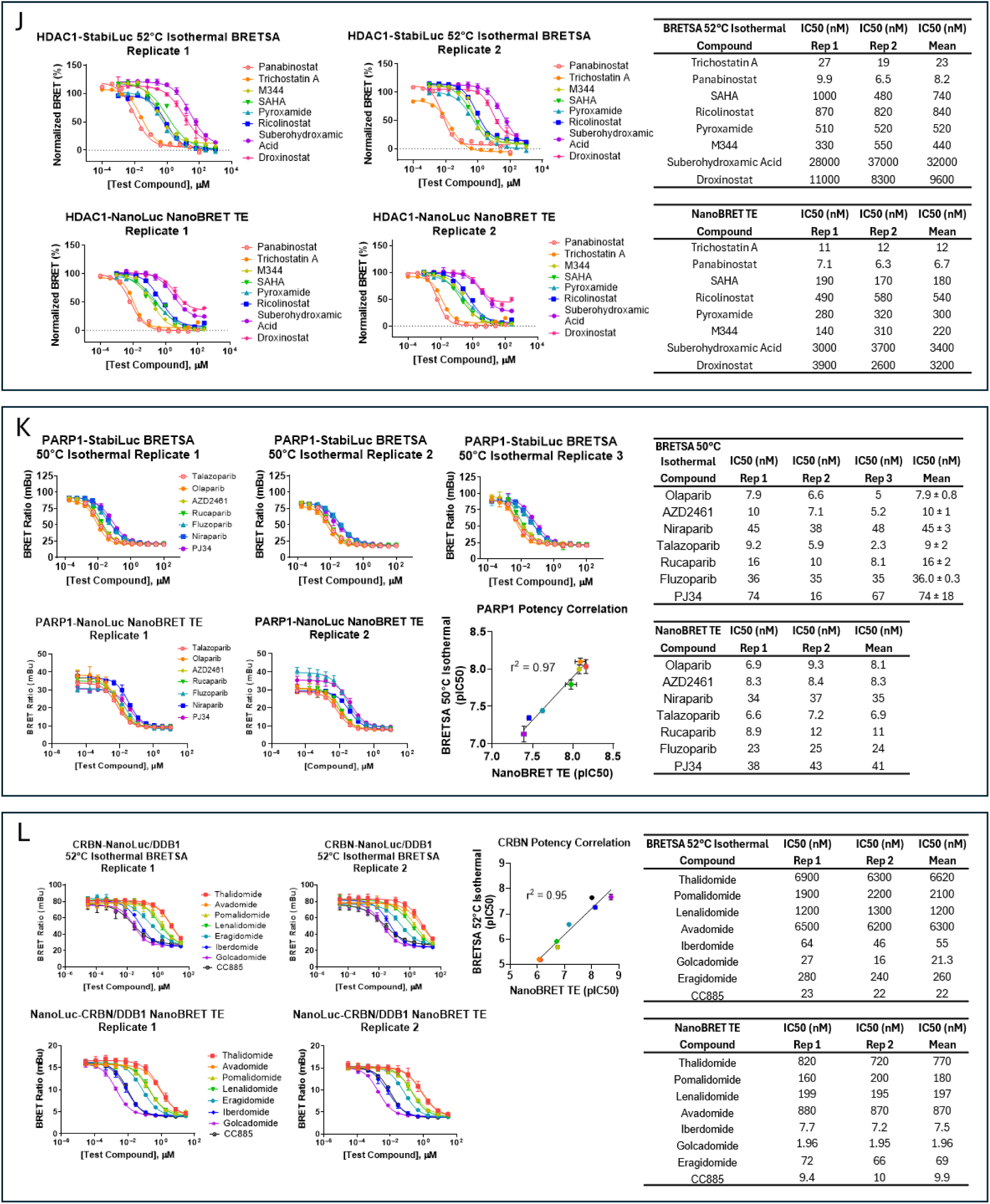
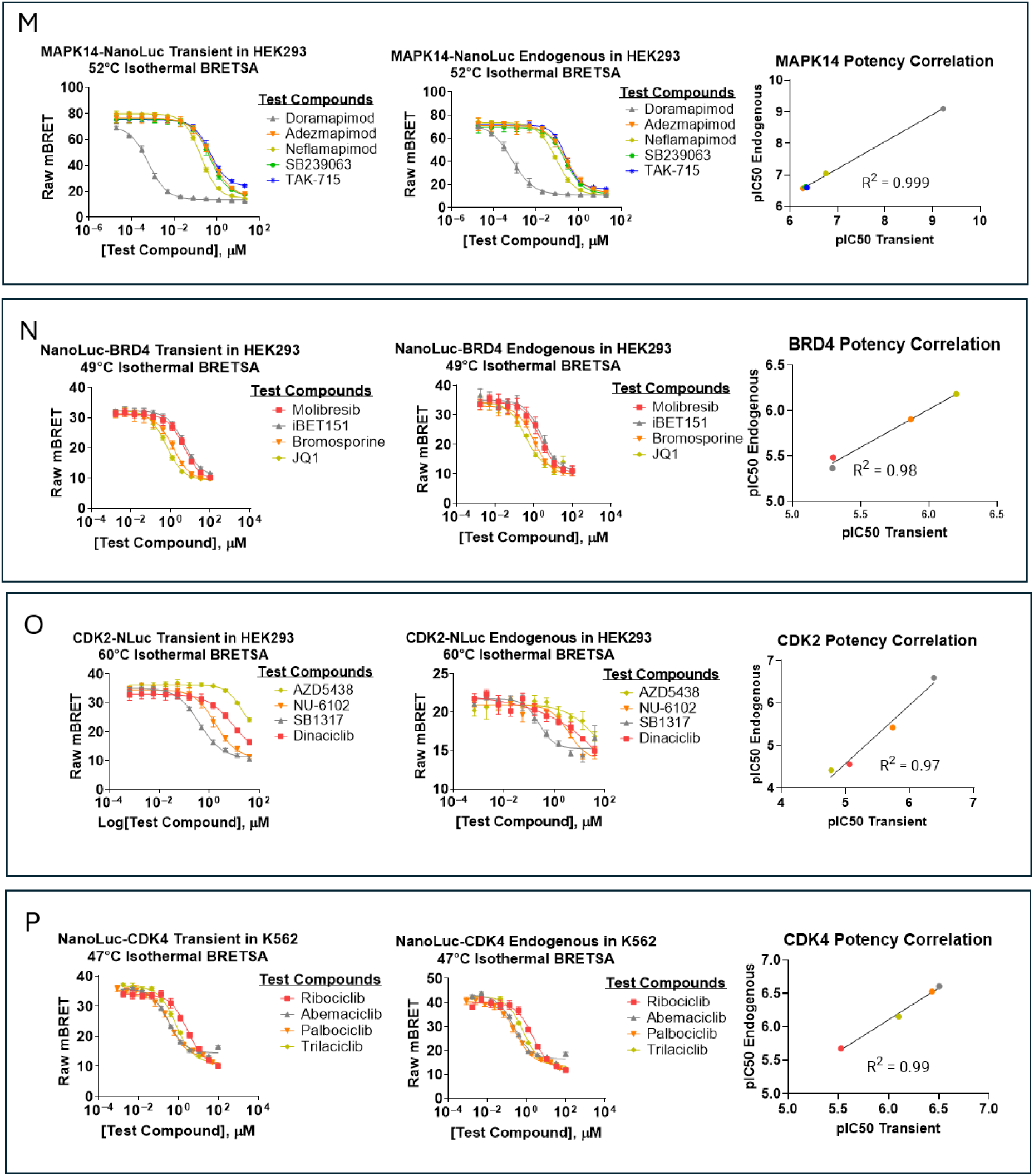
Isothermal BRETSA for potency determination. Related to Fig. 4. (A–F) BRETSA Isothermal potency profiles for representative drug targets demonstrating clear stratification of the ligand series including MAPK14 (A), AURKB (B), ABL1 (C), HSP90AB1 (D), USP7 (207-650) (E), and KRAS(G12C) (F). Compound incubation times for all targets were 1 hr, with the exception of KRAS(G12C) which used 2 hr. Data are from two independent experiments, where individual data points represent the mean (± SD) of 3–4 technical replicates. Tables with IC50 values for each replicate and the corresponding mean of the two replicates are presented in each panel (A–F). (G) Biological replicates for temperature dependent isothermal potency of reversible and irreversible covalent KRAS(G12C) SII-P ligands. Data are from two independent experiments, where individual data points are the mean (± range) of technical duplicates. (H) Biological replicates of 70°C isothermal potency of sotorasib across RAS WT and G12C alleles. Data are from two independent experiments where each individual data point is the mean (± range) of technical duplicates. Complete melt data sets are available in **Supplementary Data Sets 1 and 2**. (I) Biological replicates for temperature dependent isothermal potency of reversible and irreversible covalent AURKA ligands. For isothermal dose response curves, data are from two independent experiments where individual data points are the mean (± range) of technical duplicates. For the IC50 vs temperature replot, data are the mean (± range) of two independent experiments. (J) HDAC1 ligand potency profiles for isothermal BRETSA vs NanoBRET TE. Data are two independent experiments, where individual data points represent the mean (± range) of technical duplicates. (K) PARP1 ligand potency profiles for isothermal BRETSA vs NanoBRET TE. For BRETSA, data are representative of experiments performed three times, where individual data points represent the mean (± SD) of 4 technical replicates. For NanoBRET TE, data are representative of experiments performed twice, where individual data points represent the mean (± range) of technical duplicates. (L) CRBN ligand potency profiles for isothermal BRETSA vs NanoBRET TE. Data are representative of experiments performed twice, where individual data points represent the mean (± SD) of 4 technical replicates. (M–P) BRETSA Isothermal potency analysis for MAPK14 (M), BRD4 (N), CDK2 (O), and CDK4 (P) in transient vs CRISPR edited endogenous formats in either HEK-293 or K562 cells, as indicated. All data points for isothermal dose response curves are the mean (± SD) from a single experiment. The data points in the pIC50 correlations are the mean from a single experiment.

**Extended Data Figure 5.**
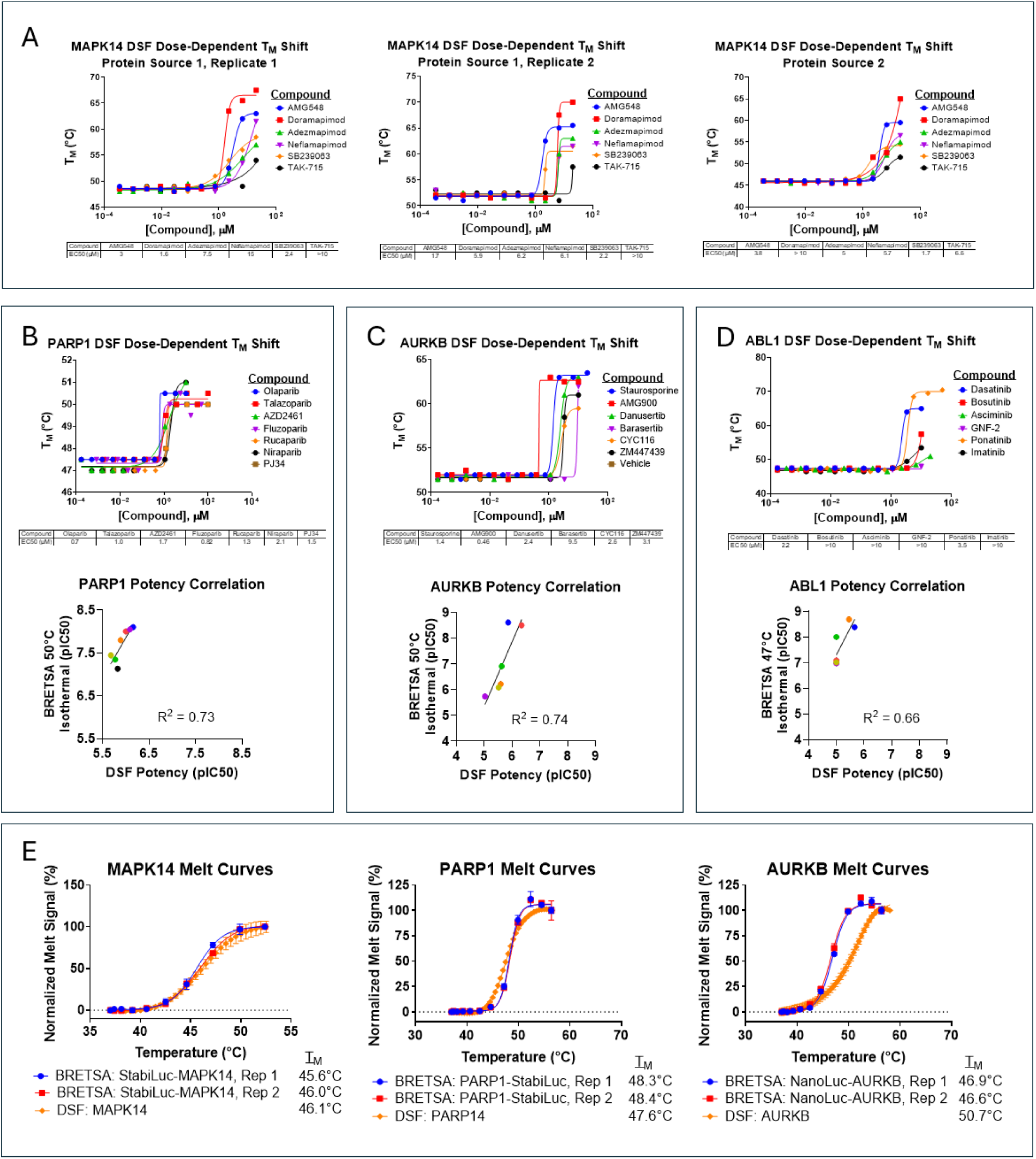
BRETSA compared to DSF. Related to Fig. 5. (A) Biological replicates of MAPK14 DSF potency profile using dose-dependent T_M_ shifts. Data are representative of experiments performed one or two times, where T_M_ values at each dose were extracted from melt curves collected in technical singlicate **[Supplementary Data Set 3]**. (B–D) DSF potency for PARP1, AURKB, and ABL1 using dose-dependent T_M_ shifts and correlation to BRETSA isothermal potencies. DSF T_M_ values at each dose were extracted from melt curves collected in technical singlicate **[Supplementary Data Set 3]**. BRETSA potency values were re-plotted from **Extended Data Figure 4**. (E) Comparison of melt temperatures observed between BRETSA and DSF for matched full length protein ORFs (MAPK14, PARP1, and AURKB). BRETSA data are replotted from **Supplementary Data Set 1** and represent two independent experiments where each data point is the mean (± range) of technical duplicates. For DSF, the data are from a single experiment where each data point is either a technical singlicate or the mean (± range) of technical duplicates **[Supplementary Data Set 3].**

**Extended Data Figure 6.**
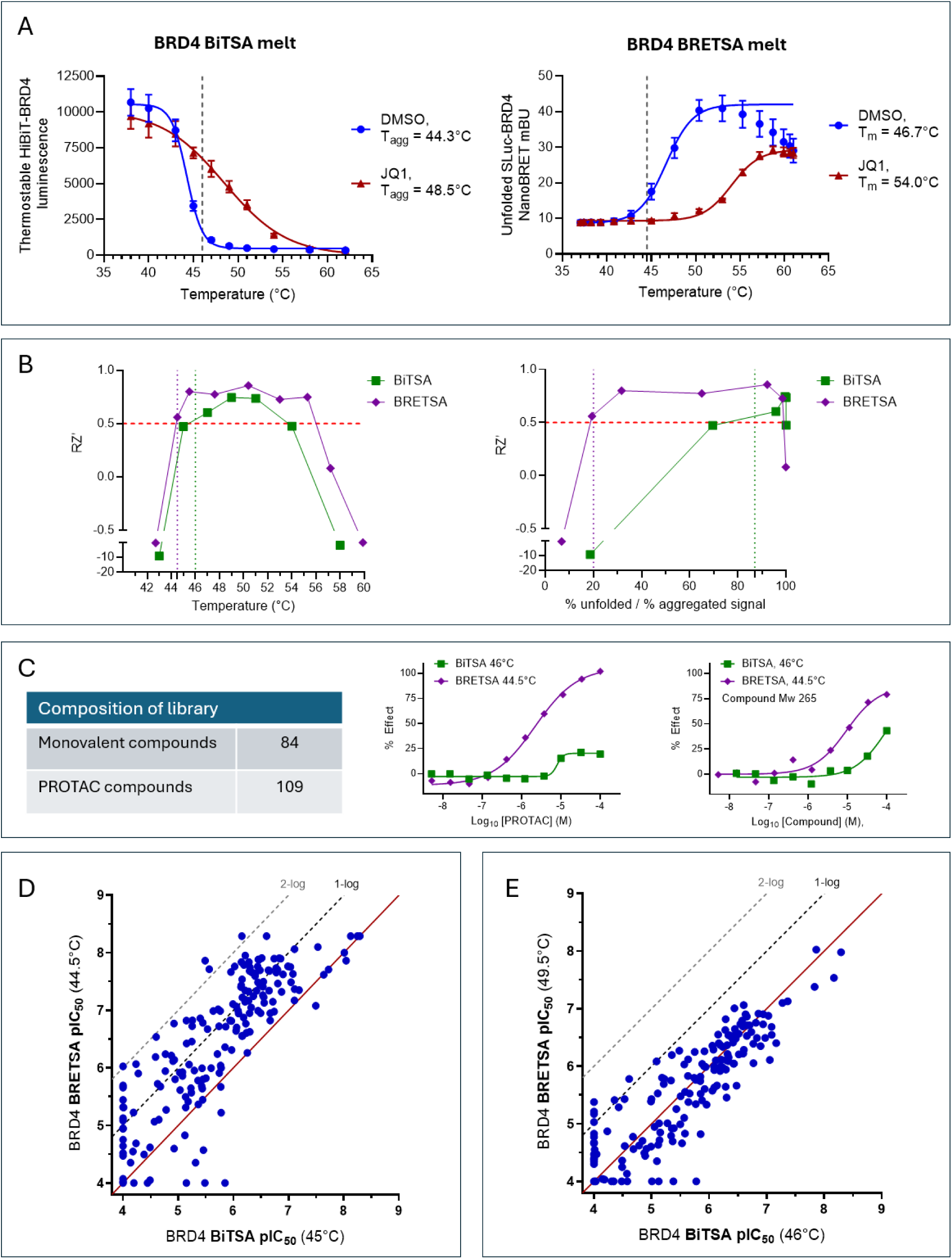
Comparison of HiBiT-BRD4 BiTSA with StabiLuc-BRD4 BRETSA. Related to Fig. 5. (A) Comparison of HiBiT-BRD4 BiTSA melt profile (data are the mean ± SD of n≥9) with StabiLuc-BRD4 BRETSA melt profile (data are the mean ± SD of n=4 across 2 replicates) in the absence or presence of 30 µM JQ1. Dashed lines indicate the lowest robust assay temperature selected for compound screening in Figure 5. (B) Assay robustness determined from Max/Min responses of the HiBiT-BRD4 BiTSA assay (Green) or the StabiLuc-BRD4 BRETSA™ assay (Purple) as a function of temperature. A robust assay temperature was determined as RZ’ >0.5. For BiTSA, this was achieved at 46°C (left panel), representing 87% aggregated BRD4 signal (right panel). For BRETSA, this was achieved at 44.5°C (left panel), representing 20% unfolded BRD4 signal (right panel). (C) Summary of the composition of a library of 193 BET-containing pharmacophores screened for IC50 in BiTSA and BRETSA across various assay temperatures with additional representative dose response curves from 46°C BiTSA and 44.5°C BRETSA for a PROTAC and a low Molecular weight (MW 265) monovalent compound. (D) Alternative BiTSA vs BRETSA assay temperature correlation plot. When BiTSA assay temperature was further reduced to 45°C, the BRETSA assay retained potency advantages. (E) Alternative BiTSA vs BRETSA assay temperature correlation plot. When BRETSA assay temperature was increased to 49.5°C to achieve 87% unfolded BRD4 signal comparable to the 87% aggregated BRD4 signal in BiTSA 46°C, compound potencies were comparable, confirming the temperature-dependence of the apparent potency advantages.

**Extended Data Figure 7.**
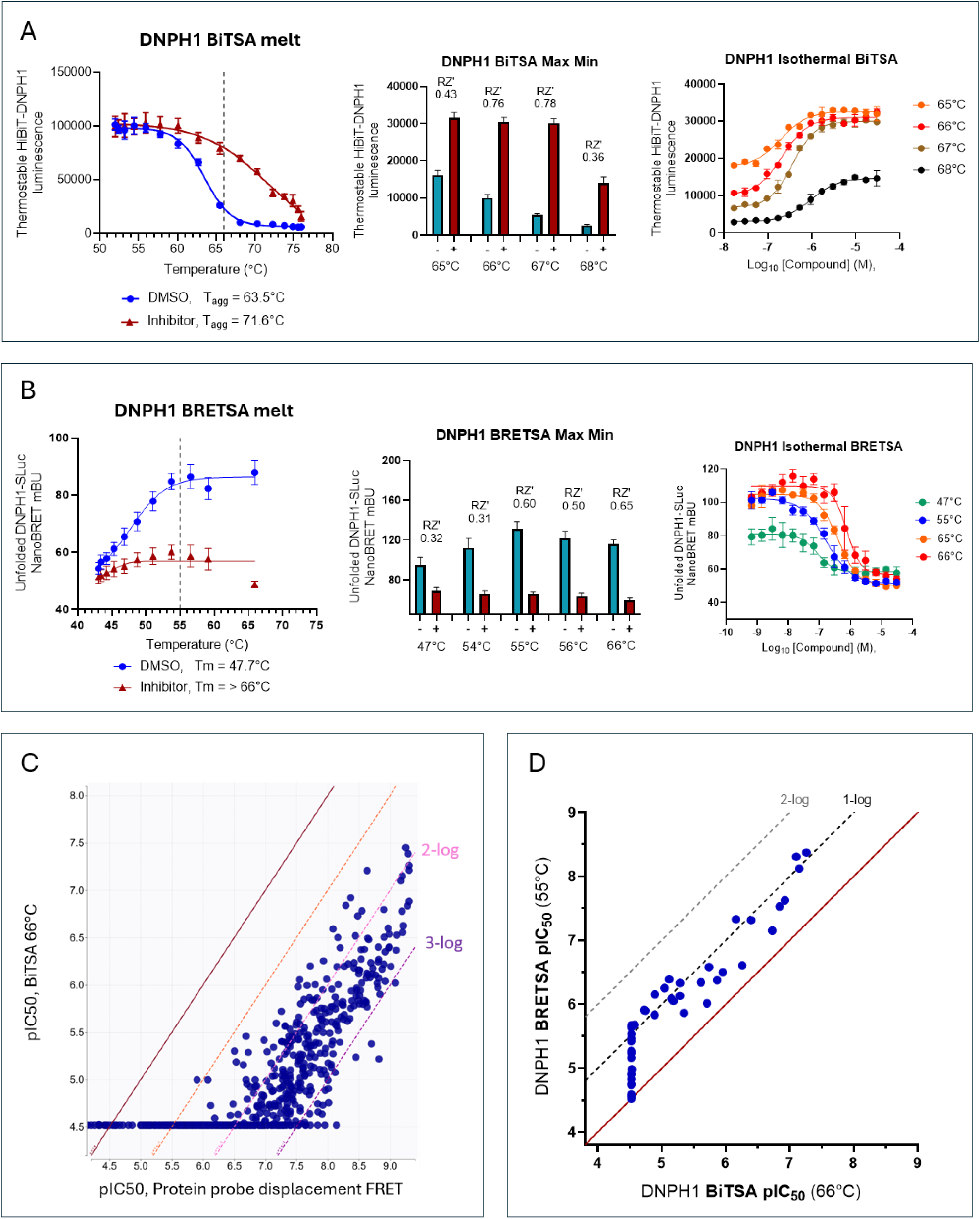
Comparison of HiBiT-DNPH1 BiTSA with DNPH1-StabiLuc BRETSA. Related to Fig. 5. (A) Determination of the lowest robust assay temperature for DNPH1 BiTSA. Left panel, HiBiT-DNPH1 BiTSA melt profile (data the mean ± SD of n=6) in the absence, or presence, of 30µM DNPH1 Inhibitor. Dashed line indicates the lowest robust assay temperature. Middle panel, assay robustness determined from Max DNPH1 inhibitor (+), Min DMSO (-) responses of the HiBiT-DNPH1 BiTSA assay as a function of temperature (data are the mean ± SD of n=16). A robust assay was determined as RZ’ >0.5. Right panel, corresponding isothermal dose response curves for a representative DNPH1 inhibitor supporting choice of 66°C as the lowest robust BiTSA temperature (data are the mean ± span of n=2). (B) Determination of the lowest robust assay temperature for DNPH1 BRETSA. Left panel, DNPH1-StabiLuc BRETSA melt profile (data are the mean ± SD of n=24) in the absence, or presence, of 30µM DNPH1 Inhibitor. Dashed line indicates the lowest robust assay temperature. Middle panel, assay robustness determined from Max DNPH1 inhibitor (+), Min DMSO (-) responses of the DNPH1-StabiLuc BRETSA assay as a function of temperature (data the mean ± SD of n=24). A robust assay was determined as RZ’ >0.5. Right panel, corresponding isothermal dose response curves for a representative DNPH1 inhibitor supporting 55°C as a robust BRETSA assay temperature (data the mean ± SD of n=3). (C) Correlation of DNPH1 66°C BiTSA pIC50 with a recombinant DNPH1 protein probe-displacement FRET binding assay for 641 DNPH1 inhibitors showing 100-to-1000-fold weaker apparent potency in BiTSA. Composition of the chemical series tested, and details of the recombinant protein binding assay are reported in *Barlaam et al., J. Med. Chem, 2025.* (D) BiTSA vs BRETSA correlation plot for the same 60 compounds reported in Figure 5F.

**Extended Data Figure 8.**
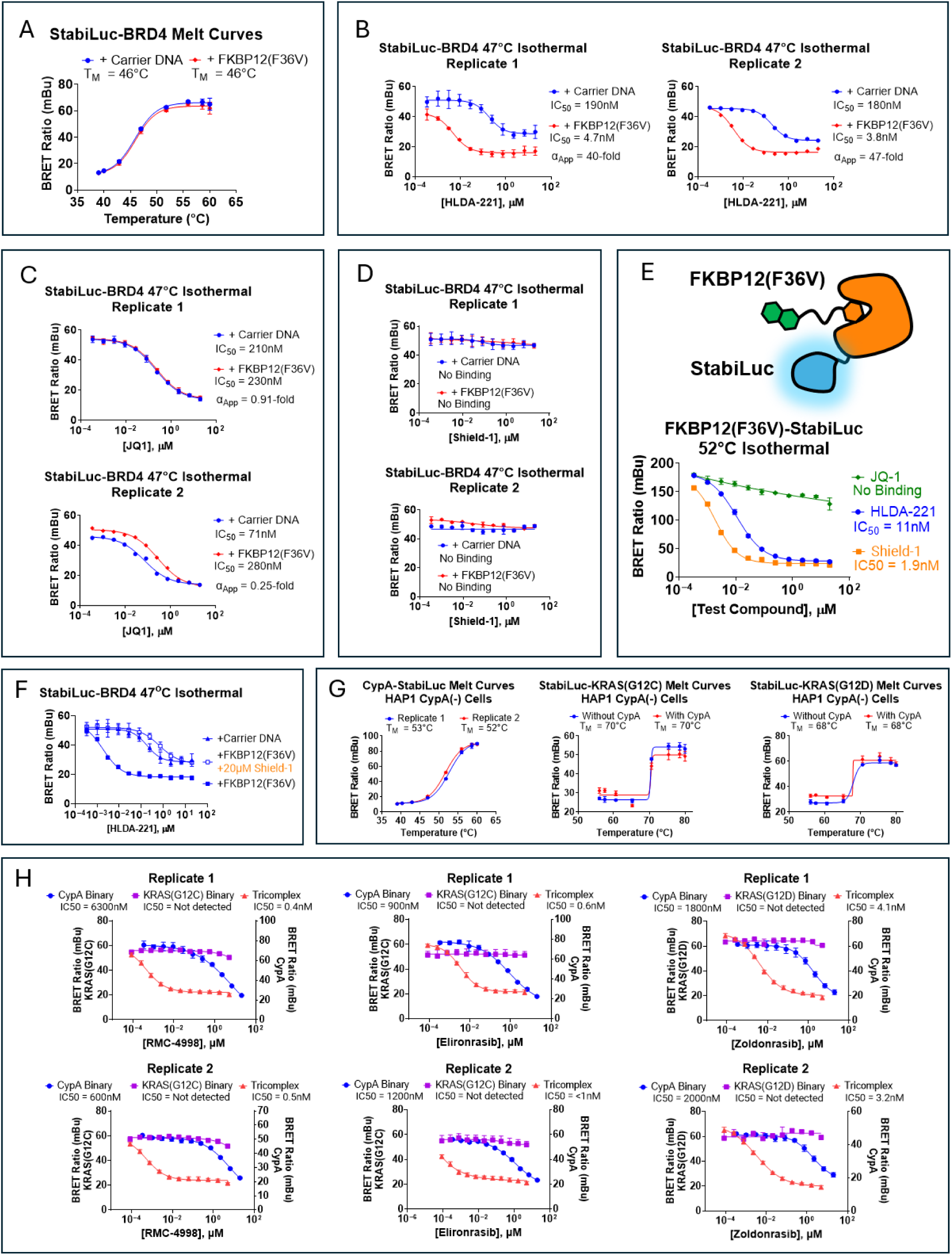
Cellular TE and cooperativity for a model RIPTAC and non-degrading molecular glues. Related to Fig. 6. (A) Raw melt curves for StabiLuc-BRD4 ± FKBP12(F36V) co-expression. (B–D) biological replicates of isothermal dose response curves for StabiLuc-BRD4 ± FKBP12(F36V) for HLDA-221 (B), JQ1 (C), and Shield-1 (D). For panel B, individual data points of replicate 1 and replicate 2 are the mean (± SD) of 4 technical replicates or a technical singlicate, respectively. For panel C, individual data points of replicate 1 and replicate 2 are the mean (± SD) of 4 technical replicates or a technical singlicate, respectively. For panel D, individual data points of replicate 1 and replicate 2 are the mean (± SD) of 4 technical replicates or the mean (± range) of technical duplicates. (E) FKBP12(F36V) isothermal potency for HLDA-221, JQ1, and Shield-1. Individual data points are the mean (± SD) of 4 technical replicates from a single experiment. (F) Reversal of BRD4 cooperativity for HLDA-221 by co-treatment with 20µM Shield-1. Data are the mean (± SD) of 4 technical replicates from a single experiment. (G) Melt curves for CypA-StabiLuc, StabiLuc-KRAS(G12C) and StabiLuc-KRAS(G12D) in HAP1 CypA(-) cells. CypA data are from two independent experiments where individual data points are the mean (± SD) of 4 technical replicates. KRAS data are from a single experiment in which melts were compared in the presence or absence of CypA co-transfection as described in the methods, where individual data points are the mean (± SD) of 4 technical replicates. (H) Biological duplicates for isothermal BRETSA potency of RMC-4998, Elironrasib, and Zoldonrasib for the CypA binary configuration, the KRAS binary configuration, or the Tricomplex configuration as described in Fig. 6f and the methods. Individual data points are the mean (± SD) of 4 technical replicates.

